# Environmental constraints can explain clutch size differences between urban and forest blue tits: insights from an egg removal experiment

**DOI:** 10.1101/2023.01.05.522710

**Authors:** Mark D. Pitt, Pablo Capilla-Lasheras, Norah S.S. Alhowiti, Claire J. Branston, Eugenio Carlon, Jelle J. Boonekamp, Davide M. Dominoni

## Abstract

Urban environments present novel ecological challenges to wild species. Understanding whether species responses to urban living are adaptive or maladaptive is critical to predicting the impacts of urbanisation on biodiversity. In birds, urban populations generally exhibit reduced reproductive investment (clutch size) compared to forest populations. However, whether smaller clutches are adaptive, or a result of environmental constraints is unclear. Here, to investigate these two hypotheses, we quantified the ability of urban and forest blue tits (*Cyanistes caeruleus*) to lay new eggs upon egg removal. Consistent with the constraint hypothesis, our results suggest that urban females do not lay new eggs, at least to the same extent as forest birds. Meanwhile, forest birds laid approximately two additional eggs. As urban blue tits did not lay replacement eggs, our experiment resulted in a brood reduction and nestlings from urban experimental nests had higher survival than those from urban control nests, suggesting that females may be misjudging urban habitat quality and produce a clutch too large to be sustained. Taken together, our results suggest that urban females may experience constraints that limit egg formation and/or exacerbate the trade-off between female survival and egg production. This has important implications for urban green space management.

## Introduction

Urban land coverage is expanding globally, increasing from 450.97 thousand km^2^ in 1990 to 747.05 thousand km^2^ in 2010, and is projected to amount to 1.6 million km^2^ by 2100 (1,2). Urban environments have a distinct set of ecological features, characterised by increased ambient temperatures (3), a high abundance of non-native species (4), increased habitat fragmentation (5), increased pollution (light, chemical, and noise) (6,7), and changes in the quality, composition, and availability of food (8–10). Urban environmental conditions create novel ecological and evolutionary pressures to which species might not be well adapted, potentially compromising the persistence of wildlife globally. Understanding adaptive and/or maladaptive biological changes associated with urbanisation is crucial to determine the current and future impact of urbanisation on species population dynamics.

Recent meta-analyses of the avian literature have suggested that among all phenotypic changes that occur in urban habitats, reductions in reproductive investment (e.g., clutch size) are pervasive and occur across a wide range of species (11–13). For example, in great tits (*Parus major*) and blue tits (*Cyanistes caeruleus*), the clutch sizes of urban breeding birds tend to be 10-20% smaller than those nesting in forests (14,15). However, it remains unclear whether the small clutches of urban birds represent a constraint imposed upon females by the urban environment when laying or reflect an adaptation to urban living.

Urban environments could impose constraints on birds during egg production, limiting the female’s ability to invest in a larger clutch. In this scenario, the clutch size birds produce would be constrained by the quality and quantity of available resources (constraint hypothesis; (16)). Egg production is metabolically demanding, with the energy required for egg-laying ranging between 13-41% above the basal metabolic rate in passerines (17). In small passerines that produce large clutches, the resources required for egg production far exceed what females can store endogenously (18,19). Therefore, birds need to source the energy and nutrients for egg formation (including proteins, antioxidants, omega-3 polyunsaturated fatty acids, and calcium), from their daily diet when laying, with invertebrates (e.g., spiders, caterpillars) being the most nutrient-rich food items (20–22). Habitat fragmentation, non-native plant species, pollution, and increased pesticide use in urban areas reduces the quality and quantity of invertebrate prey (9,10). Urban birds may attempt to compensate for the reduced availability of invertebrates by exploiting the abundant and predictable human-provisioned food from refuse and bird feeders (23). However, anthropogenic food, despite being energy-rich, is nutritionally poor and contains limited proteins, antioxidants, and omega-3 polyunsaturated fatty acids (7,24). Thus, nutrient-constrained urban birds may be already laying at their maximum, being unable to source sufficient resources to invest in forming additional eggs resulting in viable offspring. Under the constraint hypothesis, the observed clutch size of urban birds would be a result of an environmental limitation that would not maximise the female’s fitness payoffs.

Alternatively, the small clutches of urban birds may be an adaptive response to urban living (i.e., genetic change in response to selection). In this scenario, rather than being constrained by the available resources when laying, urban birds would produce a clutch size that maximises their fitness payoffs, with any deviation from this observed clutch size resulting in fewer offspring being recruited into the population (adaptive hypothesis; (25,26)). Previous research reveals that urban birds have smaller broods and fledge fewer offspring than their forest counterparts (11). The small clutch sizes of urban birds could be an adaptive response to match the number of young the parents can adequately provision (27). Small clutch sizes could allow urban parents to invest more resources into fewer nestlings and reduce sibling competition, thereby maximising the number of offspring recruited (12).

To determine whether clutch size is constrained by the environment when laying or is adapted to maximise the number of offspring recruited, previous studies outside urban ecology and evolution have undertaken egg removal experiments and observed the re-laying ability of females (16,28–30). These experiments reveal that certain species can replace eggs in response to egg removal (known as indeterminate layers) (31). In indeterminate layers, if the observed clutch size is not constrained by environmental factors, females should lay replacement eggs following egg removal to match the optimal clutch size for the habitat (29,32). Alternatively, if egg production is constrained during laying, then females should produce fewer or no additional eggs following egg removal manipulation; if replacement eggs are produced, their size could be reduced (16,28).

Here, we investigate whether differences in clutch size between an urban and forest population of blue tits are explained by environmental constraints or if these differences could represent an adaptive strategy. Blue tits evolved as a cavity-nesting forest species but have readily colonised urban environments (33). Due to their prevalence in urban areas, willingness to use nest boxes, and ability to tolerate human disturbance, blue tits are an ideal study system for investigating how urbanisation influences reproductive decisions. Blue tits are indeterminate layers, tending to lay replacement eggs in response to egg removal (33,34). Thus, the number of eggs that females produce can be experimentally manipulated by removing eggs from the clutch before the female initiates incubation. Previous research on great tits (*Parus major*) reveals females lay two replacement eggs following the removal of the first four eggs (16,28). Blue tits are close relatives of great tits and, if they are not immediately constrained, should exhibit a similar response to egg removal (31,32).

During one breeding season, we removed the first four eggs laid in urban and forest blue tit nests and observed how this manipulation affected the number and size of eggs laid. Subsequently, we examined how egg removal influenced offspring quality at hatching, the number of successful fledglings, and fledging probability. If differences in clutch size between habitats are caused by constraints on egg production, then we predict that, following the egg removal manipulation, urban females should lay fewer and smaller additional eggs (and, therefore, produce smaller hatchlings) than forest females. If this was the case, however, nestling body mass on days six and 12 after hatching and fledging probability should be higher in experimental nests than in control nests because the resulting reduction in brood size should lead to lower intra-brood competition. If egg production in urban females is not constrained (and possibly reflects an adaptation to urban conditions), we predict that urban and forest females should lay a similar number of additional eggs of equal size following the egg removal manipulation. We would also predict no difference in hatchling body mass, fledging probability, and nestling body mass between treatment groups.

## Methods

### Study populations

We monitored one urban and one forest nest-box population of blue tits in Scotland from 1^st^ April to 30^th^ June (nest-box: made with woodcrete, 260H × 170W × 180D, hole diameter: 32mm, Schwegler, Germany). The urban population resides in a city centre park in Glasgow (Kelvingrove Park, N=28 nest-boxes included in the current study, coordinates = [55.869, −4.2851]). Kelvingrove park contains open land, trees, and small shrubs, with 40% of the tree species being non-native. The forest population resides in an ancient native oak woodland, on the east shores of Loch Lomond, 40km north of Glasgow (Scottish Centre for Ecology and the Natural Environment (SCENE), N=33 nests included in the current study, coordinates = [56.129,−4.6145]). The dominant tree species at SCENE are native broadleaved trees, with only 2% of the trees comprising non-native species (23,35).

### Experimental design

#### Assignment of treatment groups

From April 1^st^ 2022, we visited nest-boxes twice weekly to monitor nest-building activity. We increased the frequency of visits to every two days once the blue tits started lining the nest-cup to ensure we accurately recorded the first egg date (FED). We visited nests after 11:00 to ensure sufficient time for the blue tit to lay (in passerines, most egg-laying occurs shortly after sunrise (36)). Within the same habitat, once a new clutch was detected it was alternatively assigned to the control or experimental group, following a 1:1 ratio to reduce any difference in phenology occurring between treatment groups. For nests included in the study, we found no difference in the FED between treatment groups (*χ*^*2*^_*df=1*_*=0*.*124, P=0*.*725*). In all nests, once a new egg was found, we marked eggs using a non-soluble marker pen, with a number corresponding to the eggs position in the laying sequence. We photographed every egg (used to calculate egg volume; details below) in both control and experimental nests, including a measuring chart in each photograph. Control nests had no eggs removed, with the photographing and marking of eggs being the only time we disturbed control females during egg production (Figure 1). In experimental nests, we removed the first four eggs from the nest on the morning each egg was laid. Eggs removed from the experimental nests were also weighed (± 0.01g) using digital scales. We did not return any of the removed eggs back to the clutches of experimental females. In the urban habitat we included 14 control and 14 experimental clutches, while in the forest we included 17 control and 16 experimental clutches, for a total of 61 clutches. Nine clutches were initially monitored but we then removed them from the experiment because the female abandoned the clutch before incubation, and we could not determine if the clutch was completed.

**Figure 1.**
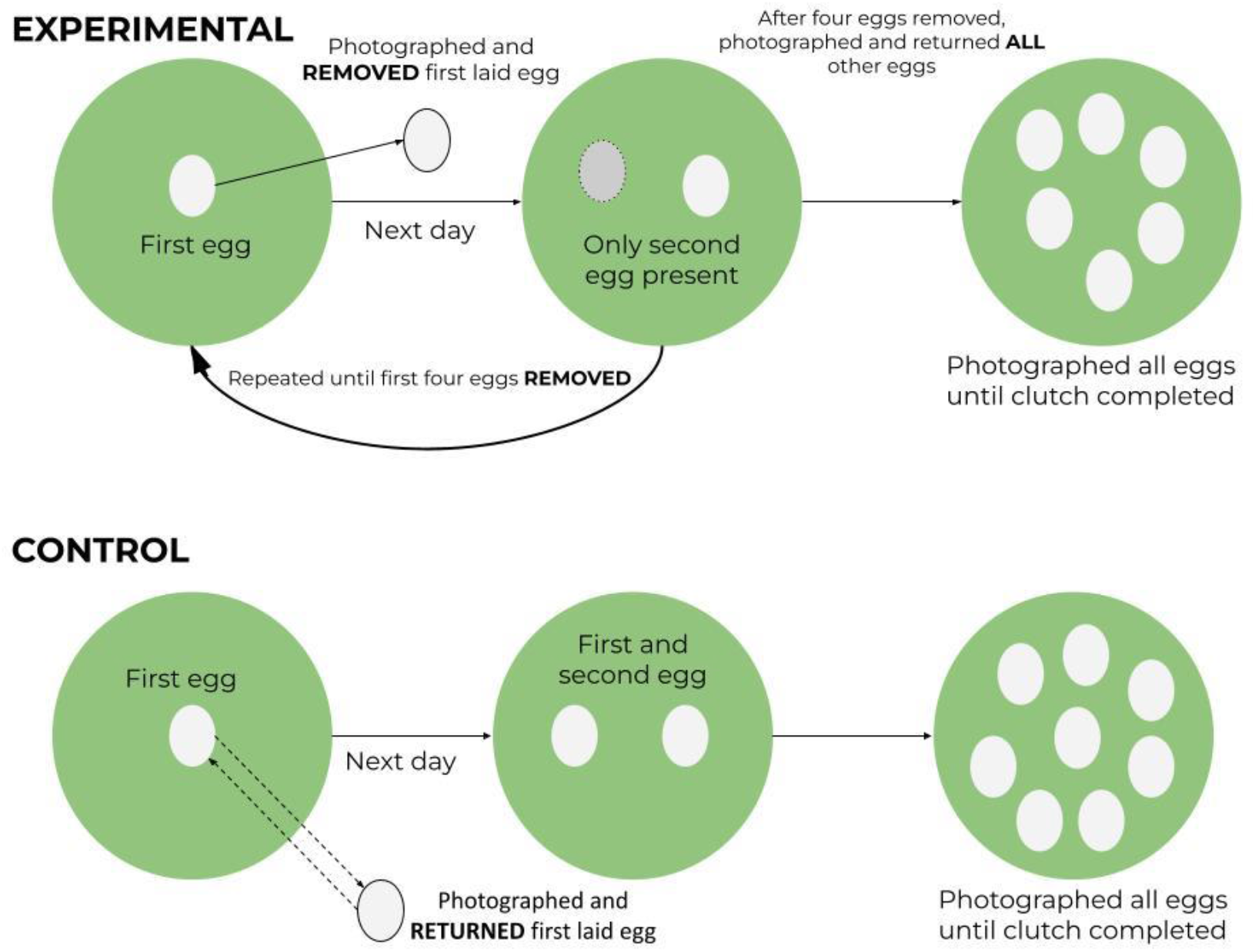
Overview of experimental design, showing the procedure for each treatment group. The first four eggs were photographed, marked, and removed from experimental nests on the day they were laid. After the fourth egg was removed, all subsequent laid eggs remained in the nest. No eggs were removed from control nests, with eggs only being handled when photographed and marked.

When the female stopped laying for two days, and the eggs were warm to touch, we assumed the clutch was complete, and incubation had commenced (37). Assuming a minimum incubation length of 14 days from clutch completion (38), 13 days after clutch completion, we started nest-box visits every two days to record the exact hatch date. As part of a different project, nestlings were cross fostered (both within and between habitats) two days after hatching. In the interest of this paper, we treat ‘habitat’ as the habitat that the nestlings were reared in, unless stated otherwise. Six days after hatching, nestlings were fitted with a British Trust for Ornithology (BTO) metal ring with a unique ID number. We also weighed nestlings to the nearest 0.1g on days two, six, and 12 after hatching. Nest-boxes were checked >21 days after hatching to identify and record dead and fledged nestlings.

#### Egg volume measurements

We used egg volume as a proxy of egg size, which reflects the level of pre-natal investment into the young before incubation and may dictate offspring quality at hatching (39). We used an Olympus TG-6 digital camera to photograph eggs in the field. We photographed eggs on a 20×20 cm measuring chart at a 90° angle to the egg’s long axis, and there were no adjustments to lens distortion. We photographed each egg four times, rotating the egg along its long axis between photographs. To calculate egg volume, we used IMAGEJ and the egg measurement plugin developed by Troscianko (40). Using the multipoint selection tool in IMAGEJ, we selected 12 anchor points around the edge of the eggs surface. For each egg, we calculated volume (mm^3^) from three separate images. All images were analysed by a single observer to minimise the risk of between-observer variation. In total, 1677 images of 559 eggs were analysed. We used the R package *rptR v*.*0*.*9*.*22* (41) to quantify the repeatability of the individual volume measurements calculated in IMAGEJ (see methods S1). The repeatability of the egg volume measurements between the three images of a single egg was low (*repeatability [95% Confidence Interval ‘CI’] = 0*.*334 [0*.*265, 0*.*421]*). However, individual females exhibited consistent egg size within clutches (*repeatability [95% CI] = 0*.*555 [0*.*446, 0*.*642]*) and mean egg volume across the three images for a single egg was correlated with egg mass (using removed eggs, N = 86 eggs, Pearson’s correlation coefficient *r [95% CI] = 0*.*719 [0*.*654, 0*.*775])*, confirming that our measurement of egg volume was capturing biologically relevant variation. In all subsequent analyses of egg volume, we used each of the three volume measurements per egg as the response variable (rather than the mean volume per egg) and included egg ID (the individual identifier for each egg) as a random effect.

### Data analysis

#### General modelling procedure

We performed all statistical analysis and data visualisation in Rstudio v.1.4.1106 (42) using R v.4.2.0 (43).We used the R package *lme4 v*.*1*.*1* (44) to build linear models and generalised and linear mixed models to explain variation in the investigated reproductive traits (see below). We initially built a global model for each reproductive trait, containing all explanatory variables and interactions with the potential to explain variation in that trait. Using the package *lmtest v*.*0*.*9* (45), we used likelihood-ratio tests (LRTs) to determine the statistical significance of each model predictor. To test the statistical significance of each term in the global model, we sequentially dropped predictors from the model and compared the model without the focal predictor against the global model using LRTs. We did not apply model simplification of single effect predictors; we only eliminated non-significant interaction effects (including quadratic terms) to ease the interpretation of single effect coefficients. In all models, date (either FED, laying date or hatching date) was expressed as the number of days since the 1^st^ of January. Date was initially fitted as both a quadratic and linear term before quadratic terms were dropped from all the models if not significant. We used the R package *performance v*.*0*.*9*.*1* (46) to assess the normality of residuals, homogeneity of variance, and the collinearity between fixed effects. We used the package *ggplot2 v*.*3*.*3*.*6* (47) to visualise the model predictions and raw data.

### Global models

#### Number of eggs laid

We used linear models to explain variation in the number of eggs laid by a female, including the FED (mean-centred) and the habitat (two-level factor, “urban” versus “forest”) × treatment group (two-level factor, “experimental” versus “control”) interaction as fixed effects. A total of 61 clutches were initially included in the analyses, with nine clutches removed from the final dataset because the female abandoned the clutch before incubation, and we could not determine if the clutch was completed. We then created two separate within-habitat models, one for the urban habitat and one for the forest habitat, including FED (mean-centred) and treatment group as fixed effects.

#### Egg volume

To determine whether there were differences in egg volume between habitats and treatment groups, we initially built a linear mixed model with egg volume as a response variable and the following explanatory variables: the total number of eggs laid by each female (mean-centred), egg-laying date (the exact day each egg was laid; mean-centred), and the interaction between habitat and treatment group. Clutch ID (a 70-level factor) and egg ID (a 460-level factor) were included as random effect intercepts. We also assessed whether the egg removal treatment differentially affected the total volume of eggs produced by a female between habitats (see methods S1; Results S1; Tables S6-S9).

#### Variation in egg volume throughout the laying sequence

We used a liner mixed model to test the effect of laying order (fitted as a continuous variable) on egg volume to determine if investment into egg quality changed over the lay sequence. We included eggs one to nine in the laying sequence in the model as only three urban birds produced a clutch larger than nine eggs. This model contained egg volume as the response variable and egg laying date (mean-centred), the total number of eggs laid by each female (mean-centred), and the three-way interaction between the position of the egg in the lay sequence (fitted as a continuous variable), habitat, and treatment group as fixed effects. We included clutch ID (a 70-level factor) and egg ID (a 460-level factor) as random effects. We ran an additional model for egg volume where laying order as fitted as a categorical predictor with two levels (eggs one to three or eggs four to nine (see methods S1; Results S2; Tables S10-S11)).

#### Hatchling and nestling body mass

We used a linear mixed model to determine whether egg removal differentially affected nestling body mass between habitats two-days after hatching. As previous research suggests there may be a relationship between egg size and hatchling body mass, the habitat or treatment group with the largest eggs should also produce heavier hatchlings. We included nestling body mass two days after hatching (the first-time nestlings were weighed) as the response variable with the number of siblings (equivalent to the number of hatched eggs; mean-centred), hatch date (mean-centred), time of day (two-level factor: “morning” or “afternoon”), and the two-way interaction between habitat and treatment group as fixed effects. Clutch ID (a 58-level factor) was included as a random effect. We also modelled nestling body mass on days six and 12 after hatching (see methods S1; Results S3; Tables S15-S17). Additionally, we tested whether nestling body mass two days after hatching predicted nestling survival (see methods S1; Results S4; Tables S18-S19).

#### Nestling survival to fledging

To investigate whether our experimental manipulation affected nesting survival to fledging, we ran two complementary analyses. Firstly, we fitted a linear mixed model with the number of alive nestlings at a given time point (i.e., nestling age, 4-level factor: “day two”, “six”, “12”, or “fledged”) as the response variable. In this model, we included, as fixed effects, treatment group, nestling age, the number of eggs incubated (mean-centred), hatch date (mean-centred), and the three-way interaction between nestling age, habitat, and treatment group. Secondly, we fitted a binomial model where the response variable represented whether an individual nestling had survived or not at a given age. The same fixed effects were included for this second model. We did not include the first day of hatching (day 0) in the analysis as blue tits hatch asynchronously. In both cases, clutch ID (a 58-level factor) was included as a random effect intercept.

## Results

### The effect of egg removal on the female’s ability to lay additional eggs

We found no differential effect of egg removal on the female’s ability to lay additional eggs between habitats (habitat × treatment group interaction, effect of egg removal on total number of eggs laid in the urban habitat ± SE= 0.517 ± 0.632, effect of egg removal on total number of eggs laid in the forest ± SE= 1.186 ± 0.854, *χ*^*2*^_*df=1*_*=2*.*069, P=0*.*150*; Table S1). However, visually, there appeared to be a difference in the ability of experimental and control females to replace the removed eggs between the urban and forest habitat (Figure 2), suggesting that the lack of statistical support of the interaction might have been due to a low sample size. The confidence intervals for the total number of eggs laid by forest females in the experimental group did not include the estimated mean of the control group and did not overlap with the confidence interval of the control group (Figure 2). In the urban site, the confidence intervals for the total number of eggs laid by females in the experimental group did include the estimated mean of the control group, and the associated confidence intervals, suggesting that females in the urban habitat did not replace removed eggs, at least to the same extent that forest females did (Figure 2). To statistically confirm our visual interpretation of the data, we then created two separate models, one for the urban and one for the forest habitat. In the urban habitat, treatment group had no effect on the number of eggs laid (effect of egg removal on total number of eggs laid ± SE=0.472 ± 0.594 eggs, *χ*^*2*^_*df=1*_*=0*.*701, P=0*.*403;* Table S2). In the forest, birds in the experimental group laid significantly more eggs than those in the control group (effect of egg removal on total number of eggs laid ± SE=1.632 ± 0.600 eggs, *χ*^*2*^_*df=1*_*=7*.*267, P=0*.*007;* Table S3).

**Figure 2.**
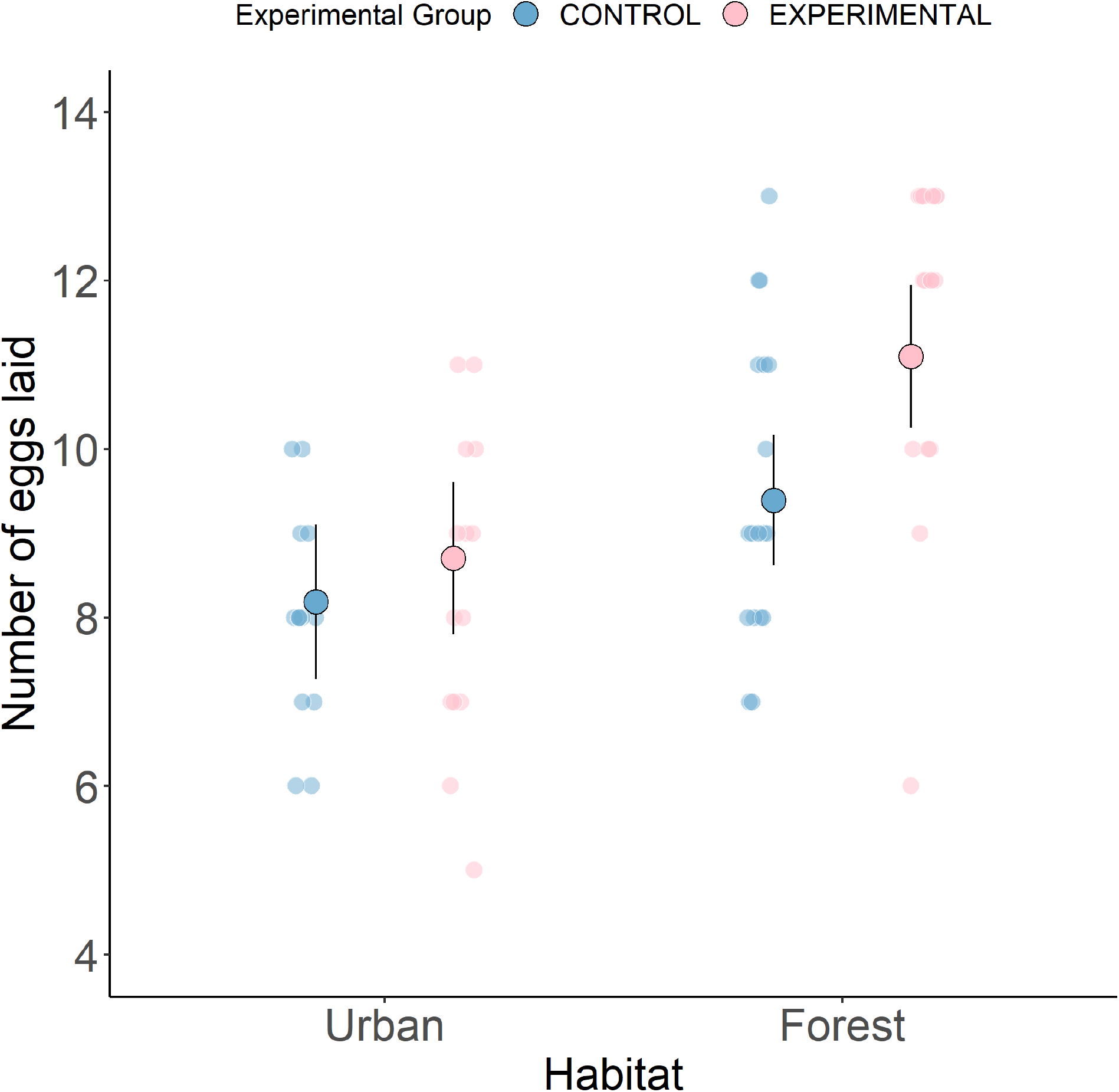
The effect of egg removal on the number of eggs laid by the females in the urban and forest habitat. Large dots represent model predictions ± 95% confidence intervals. Small dots are the raw data points. N=61 clutches.

### The effect of egg removal on egg volume

When laying order was not included in the model, variation in egg volume was not explained by the treatment group × habitat interaction (*χ*^*2*^_*df=1*_*=1*.*288, P=0*.*525;* Table S4). Additionally, when fitted as single effects (in contrast to when they were included in the interaction), habitat (*χ*^*2*^_*df=1*_*=3*.*745, P=0*.*053;* Table S5, Figure 3a) and treatment group (*χ* ^*2*^_*df=1*_*=0*.*029, P=0*.*864;* Table S5, Figure 3a) did not affect egg volume. In contrast, when laying order was included as a continuous predictor of egg volume, the three-way interaction between laying order, treatment group, and habitat significantly explained variation in egg volume (*χ*^*2*^_*df=1*_*=3*.*949, P=0*.*047;* Table S12, Figure 3b). There was no difference in egg volume across the lay sequence and between treatment groups in the urban habitat, the same held for control nests in the forest habitat. Meanwhile, egg volume declined over the lay sequence in experimental nests from the forest habitat (Figure 3b).

**Figure 3.**
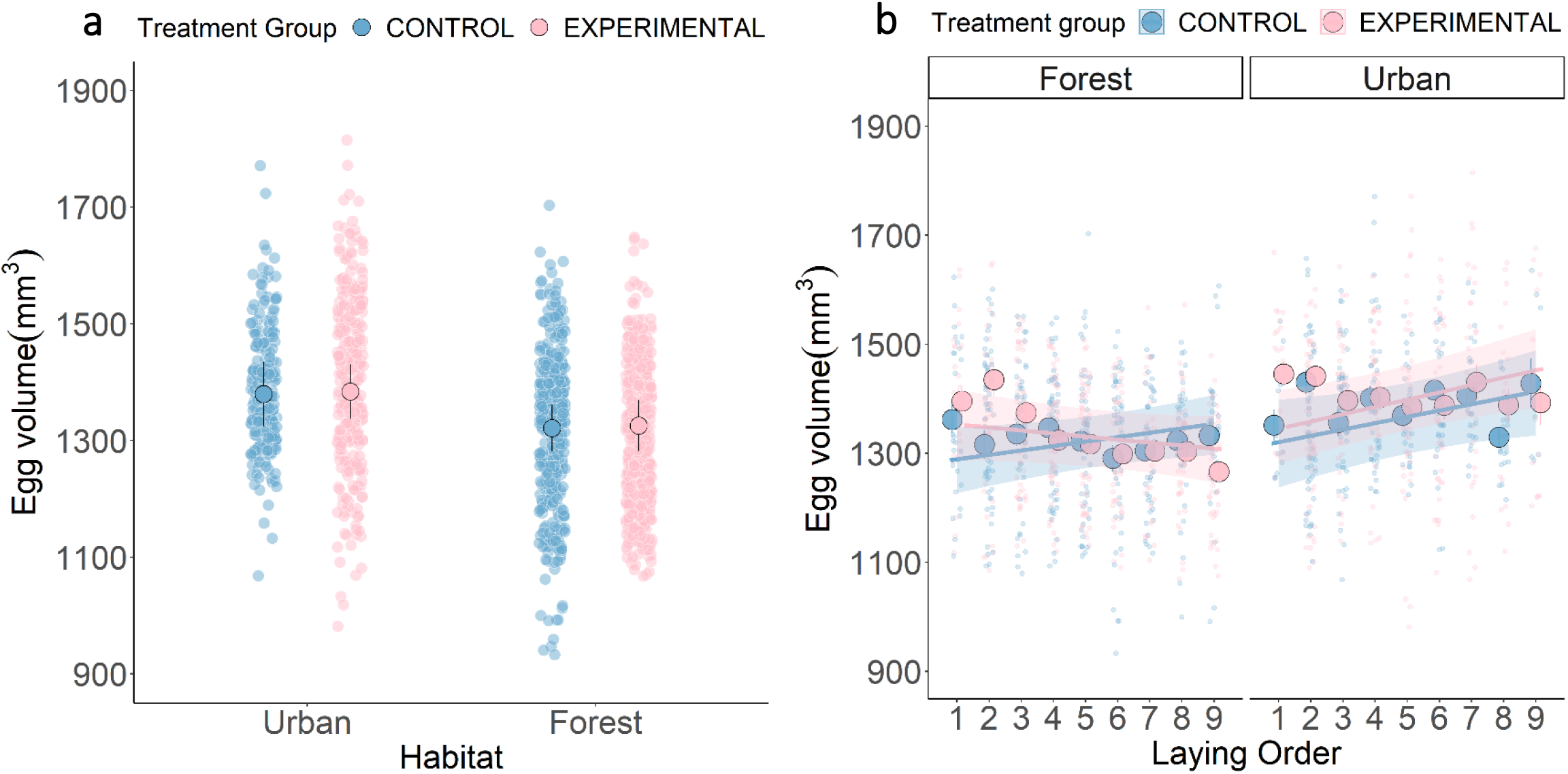
The effect of experimental egg removal on egg volume. **a)** The effect of the treatment group on egg volume in the urban and forest habitat. Large dots represent model predictions from the minimal model ± 95% confidence intervals. **b)** The effect of the treatment group and laying order (eggs one to nine) on egg volume in the urban and forest habitat. Large dots represent raw means ±se. Solid lines and shaded area represent model predictions ± 95% confidence intervals, respectively. Small dots are the raw data points. N=1380 volume measurements in both figures.

### The effect of egg removal on nestling body mass

There was no differential effect between habitats of the egg removal treatment on nestling body mass two days after hatching (treatment group × habitat interaction, *χ*^*2*^_*df=1*_*=0*.*132, P=0*.*717;* Table S13). However, we found that nestlings in experimental nests were smaller than those in control nests two days after hatching in both habitats (*χ*^*2*^_*df=1*_*=4*.*117, P=0*.*042;* Table S14, Figure 4). Nestling hatch date did not affect body mass (*χ*^*2*^_*df=1*_*=0*.*182, P=0*.*670;* Table S14). There were no differences in hatch date between habitats (*χ*^*2*^_*df=1*_*=0*.*190, P=0*.*663*) and between treatment groups (*χ*^*2*^_*df=1*_*=0*.*180, P=0*.*184*). There was no difference in day 12 body mass between forest treatment groups, but nestlings from urban experimental nests were heavier than those from urban control nests 12 days after hatching (*χ*^*2*^_*df=1*_*=1*.*746, P<0*.*001;* Table S17).

**Figure 4.**
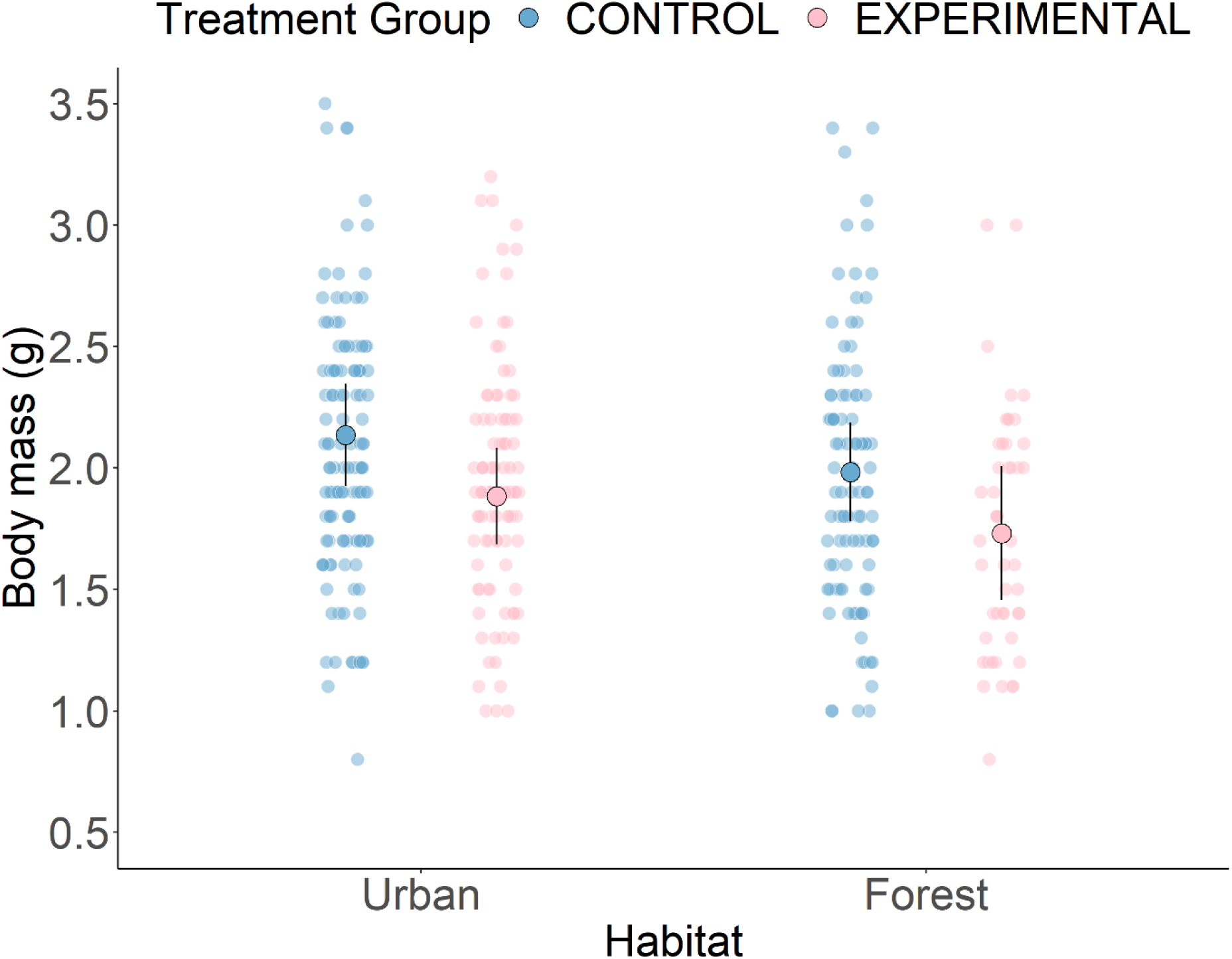
The effect of the habitat and treatment group on nestling mass (g) two days after hatching. Large dots represent minimal model predictions ± 95% confidence intervals. Small dots are the raw data points. N=372 nestlings.

### The effect of egg removal on nestling survival between habitats

We found that, while control broods in the forest were larger than experimental broods at each measured time point, in the urban habitat, control broods were larger than experimental broods only two and six days after hatching. On day 12 and at fledging, there was no difference between treatment groups in the number of nestlings alive in the urban habitat (treatment group x habitat x nestling age, *χ*^*2*^_*df=3*_*=19*.*404, P<0*.*001;* Table S20, Figure 5a). Additionally, there was no difference in nestling survival probability between treatment groups in the forest at each measured time point. Likewise, in the urban habitat there was no difference in survival probability between treatment groups on days two and six after hatching. However, on day 12 and at fledging, nestling survival was higher in experimental nests than in control nests in the urban habitat (treatment group x habitat x nestling age, *χ*^*2*^_*df=3*_*=8*.*761, P=0*.*033;* Table S21, Figure 5b).

**Figure 5.**
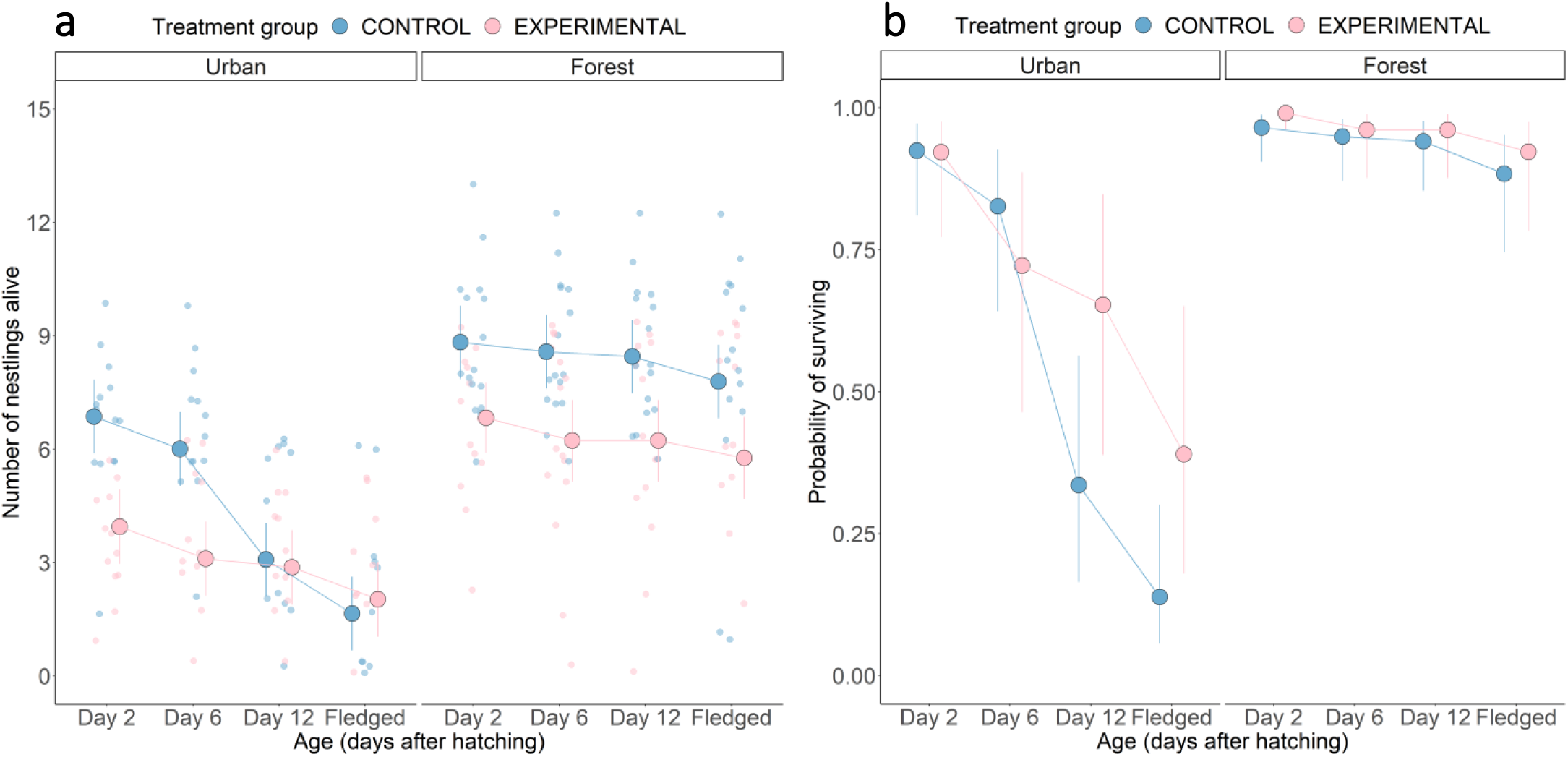
The effect of experimental egg removal on nestling survival. **a)** The effect of treatment group, habitat, and age on the number of nestlings alive. Large dots and associated bars represent model predictions ± 95% confidence intervals. Small dots represent raw data points. **b)** The effect of treatment group, habitat, and age on the probability of survival. Large dots represent model predictions ± 95% confidence intervals. N=60 broods in both cases.

## Discussion

Urban birds consistently show reduced investment to their clutch size compared to their forest counterparts (11–13), but whether this is a constraint or adaptation to the urban environment has remained unclear. To test if the smaller clutches of urban birds reflected constraints imposed upon females by the urban environment or whether this reduction might represent an adaptation to urban environments, we investigated the ability of females to replace eggs following egg removal in both urban and forest habitats. In line with the constraint hypothesis, our findings suggest that urban birds did not replace the removed eggs to the same extent that forest females did. If urban blue tits were not constrained in some way during egg production, then, as observed in the forest habitat, urban females should have laid additional eggs to compensate for egg removal and maintain their clutch size (16,28,32). This result needs to be cautiously interpreted as evidence came from habitat-specific models. However, our findings based on the experimental effect size across habitats represent the first evidence for the mechanisms behind the differences in avian clutch size between urban and non-urban habitats. There was no difference in egg volume over the lay sequence between urban treatment groups, but egg volume declined over the lay sequence in experimental forest nests. Nestlings from experimental nests also had a smaller body mass than those in control nests in both habitats two days after hatching. Additionally, due to urban birds not laying additional eggs after egg removal, this created an unintentional artificial brood reduction in the city. Urban nestlings in experimental nests were heavier at day 12 and more likely to fledge than their control counterparts, although the total number of nestlings that fledged did not differ between urban control and experimental nests, despite larger clutch sizes in the control nests. Therefore, urban birds could be producing clutches that are too large to be sustained in the urban habitat. Overall, these results suggest the environmental forces operating on clutch size differ between the urban and forest habitat. Urban conditions likely impose constraints on females when laying, preventing increased investment in egg production, such as energy and nutrient limitations (11,16). Alternatively, the smaller clutches of urban birds could be an adaptation to maximise the mother’s survival if producing large clutches incurs a survival cost to the mother (i.e., sacrificing somatic maintenance or future fecundity) (30,50). We discuss these alternative explanations of our findings below, including the limitations of this research and future directions to further understand the evolution of clutch size in anthropogenic environments.

### The effects of energy and nutrient limitation on egg production

The energetic costs associated with egg production could be increased in urban areas, preventing urban birds from replacing eggs after egg removal. As the environment and resources available to urban birds could be of low quality, urban females may differentially allocate energy between activities compared to forest birds. During the pre-laying period, individuals divide energy between nutrient acquisition, thermoregulation, territory defence, mate attraction, and egg production (51). Individuals cannot maximise their investment in all tasks, i.e., trade-offs occur (52). For example, the energy costs associated with thermoregulation may limit the energy allocated to egg production (53). In great tits, the ability to replace eggs following egg removal is temperature constrained, with females failing to upregulate egg production in years with lower spring temperatures (28). Low temperatures increase the daily energy expenditure of females during egg laying as they expend more energy on thermoregulation (54). However, in our study it is unlikely temperature prevented urban birds from replacing eggs. Urban areas are generally 1-3°C warmer than surrounding natural areas due to the heat-island effect (3). Thus, urban birds should spend less energy thermoregulating than forest birds due to the urban heat-island. If anything, urban birds should have increased the energy allocated to egg production and laid replacement eggs.

Alternatively, urban blue tits may not upregulate egg production after egg removal due to being constrained by the poor nutrient quality of anthropogenic food (constraint hypothesis). In small passerines producing large clutches, the reserves required for egg production exceed what females can store endogenously (18). Thus, they must acquire the nutrients for egg production daily from their diet when laying (19). Following egg removal, urban females may have been unable to source sufficient high-quality resources to invest in the production of additional eggs. Previous results suggesting that the diet of urban blue tits is of lower quality that the diet of forest blue tits are in line with this explanation (23).

Given females deplete endogenous biomolecules by reallocating them to the egg, the mother may face a trade-off between allocating resources for somatic maintenance and egg production, which could underlie the costs of egg production (28,30). During laying, females have increased susceptibility to oxidative stress (55), disease (56), and parasite infection (57). The effects of nutrient limitation during egg laying on female condition and survival may have been operating differently between habitats. On one hand, the nutrient requirements for the female’s somatic maintenance may be fulfilled in the forest habitat. Thus, forest females may invest any additional resources into upregulating egg production at little cost to their survival. On the other hand, as nutrient limitation may be higher in urban areas (7,23,27), urban females in poor condition could be selectively investing in somatic maintenance at the expense of egg quantity (24,58). Thus, after egg removal, urban females may have insufficient nutrient reserves to form additional eggs, as doing so would reduce survival or future fecundity (16,50).

### The trade-off between clutch size and egg volume

As the resources available to the breeding female are limited, individuals producing larger clutches may compensate by producing smaller eggs (59). However, we found that egg volume did not differ between habitats, despite urban birds producing smaller clutches than their forest counterparts. For both treatment groups in the urban habitat, we found that egg volume exhibited no change over the lay sequence: the same holds for control clutches in the forest habitat. Despite this, previous studies in small passerines found egg size increased over the lay sequence, reflecting increased protein content in the last laid eggs (16,39), which may compensate for the detrimental effects of asynchronous hatching (60). In our experimental forest nests, egg volume declined over the lay sequence. These results are in line with previous studies finding females have a limited ability to maintain egg quality when laying above their usual clutch size (51). If urban birds laid replacement eggs, this could detrimentally impact egg and offspring quality, which they may be unable to compensate for during nestling rearing.

Thus, in urban areas, females may terminate laying earlier than forest birds before they pass their critical physiological threshold where egg size and clutch size are traded off against each other. Meanwhile, forest blue tits might be able to upregulate egg production following egg removal, as they can compensate for any detrimental effects of egg size on offspring quality later in the reproductive cycle.

### Egg size and nestling mass at hatching

As parents can directly affect their nestling’s body mass by maximising investment in egg quality (39), one may predict that an increase in egg size may be advantageous, especially in urban areas where resources are limited in the nestling-rearing phase. Large eggs should result in nestlings with increased nutrient reserves, facilitating rapid growth and boosting immunocompetence after hatching (48,61). We found no difference in nestling body mass two days after hatching between the urban and forest habitats, but experimental nestlings were smaller than control nestlings in both habitats. Hatchlings in experimental nests may be smaller than those in control nests because they are on average younger. Blue tits initiate incubation before clutch completion, with the first laid eggs hatching *en masse* while later laid eggs hatch over a more prolonged period (33). As we aged nestlings from the day the first nestling hatched, the average age of nestlings in control nests may be higher than those in experimental nests, who come from eggs later in the lay sequence and may have only just hatched.

### Maladaptive responses to the urban environment

In the urban habitat, nestling survival was greater in experimental nests, where parents reared fewer nestlings than in control nests. This was a by-product of our experimental manipulation as urban birds did not re-lay and we did not return the removed eggs back to the nest, which resulted in a brood reduction. This result might suggest that urban females could be making a maladaptive decision when laying as they seemingly lay more eggs than the number of nestlings they can successfully fledge. The clutches of urban birds may be too large due to the lack of barriers preventing gene flow from neighbouring forests (62,63). Immigration between the urban and forest habitat may have homogenised gene pools, preventing the evolution of adaptive clutch size if the two populations have a different fitness optimum for clutch size (64). Alternatively, urban environments may function as ecological traps (7). Although urban areas may have more resources and stable conditions during the winter than forest habitats, leading birds to preferentially settle in cities, the poor food quality and exposure to pollution may impair nestling development and survival to fledging (23,65). Therefore, urban birds may be misinterpreting habitat quality when settling and, subsequently, produce clutches that are too large to be sustained (66).

### Limitations and future investigations

This study has limitations that could influence the interpretation of the results. First, most females were not individually identified. We were, therefore, unable to assess how parental quality and age affected the ability of females to respond to our experimental manipulation. Low-quality or young individuals that naturally produce smaller clutches may be pushed into urban areas if free territories in the forest are no longer available (7). Indeed, laying following egg removal may be more common in forest habitats if females are more experienced at breeding (57,67). Therefore, in the future, we need to determine if the parental quality or the age structure of the urban and forest populations differ and if that further explains differences in clutch size between urban and forest habitats. Second, this study consists of only one urban and one forest population, in one year. Replicated work is needed to assess if our results generalise across study populations, habitats, and temporally. Species may also exhibit different responses to egg removal if the factor limiting egg production varies between species or between years. Information on the costs of egg production should be obtained from urban and natural populations of different species filling various niches. Third, we appreciate that number of fledglings is only a proxy for fitness that only partly captures the number of offspring recruited across the parent’s lifetime. Fourth, future work should compare the costs of each stage of the reproductive cycle (egg laying, incubation, and nestling rearing) between urban and forest birds, and assess the consequences of these costs, as there may be interactive effects between reproductive phases that operate differently between urban and forest habitats (29). Finally, a common garden or transplant experiment would be valuable to further consolidate our findings and confirm that the small clutches of urban birds reflect an environmental constraint.

## Conclusion

Our study provides initial experimental support for blue tits being more constrained during egg production in urban than forest habitats. Urban birds may experience greater energetic or nutrient constraints than forest birds that either immediately restricts egg formation or exacerbates the trade-off between somatic maintenance and egg production in urban breeding females. Additionally, urban birds may produce small clutches as an increase in clutch size would be traded off against egg size, the detrimental effects of which city birds cannot compensate for during the nestling rearing stage. Our results also suggest urban birds produce clutches too large to be sustained. Egg removal in urban experimental nests resulted in a brood reduction, and the nestlings in the reduced broods had higher survival prospects than those in control nests. Thus, the clutch sizes of urban birds may be maladaptive, either due to gene flow between the forest and the city preventing the evolution of clutch size or through birds misjudging the quality of the urban environment. Overall, our results emphasise a need to incorporate the environmental constraints and fitness costs associated with egg production when attempting to explain variation in reproductive investment and success for birds breeding in anthropogenically modified landscapes.

## Ethics Statement

All work involving nest disturbance, egg removal, and cross-fostering was covered by the license 207317 issued by NatureScot to DMD. Permission for bird ringing was granted by the British Trust for Ornithology, with licenses to DMD (permit number: C6822), and CJB (permit number: C6271).

## Supplementary Information

Supplementary material is available online in the associated supplement files.

## Acknowledgements

We would like to thank Martha Hayward, Rachel McConnel, Rachel Reid, and Lotta Ruha for helping to monitor the nest-boxes and collect bird breeding data. We also thank the Loch Lomond and the Trossachs National Park and Glasgow City Council for allowing us access to the study areas and supporting our research. We thank the Scottish Centre for Ecology and the Natural Environment for facilitating our research.

