## Supplementary materials for "Environmental constraints can explain clutch size differences between urban and forest blue tits: insights from an egg removal experiment"

### Supplementary material

***Methods S1.***

*Repeatability of the egg volume measurements*

We used the R package *rptR v.0.9.22* (Stoffel *et al.*, 2017) to quantify the repeatability of the volume measurements calculated in IMAGEJ. To estimate repeatability, we used a mixed effects model framework, including habitat as a fixed effect. Clutch ID (a 70-level factor) and the unique identifier given to each egg (egg ID: a 559-level factor) were included as random effect intercepts. Here, repeatability was calculated for egg ID (consistency of the volume measurements per egg) and clutch ID (consistency of the volume measurements per clutch [i.e., nest-box], equivalent to how consistent egg size was within an individual female). For experimental nests, where the first four eggs laid were weighed, we used Pearson's correlation coefficient *r* to test the strength of the relationship between egg mass and egg volume.

*Total volume of egg material produced by a female*

We calculated the total egg volume produced by a female by taking the mean egg volume per nest box and multiplying it by the number of eggs laid. We then used a linear model to analyse the relationship between the total egg volume produced and the following explanatory variables: first egg laying date (mean-centred), and the interaction between habitat and treatment group. Finally, we created within-habitat models for the urban and forest habitat, with the total egg volume produced as the response variable and the first egg-laying date (mean-centred) and treatment group as fixed effects.

*The effect of egg laying group on egg volume*

We first divided eggs into two laying order groups: eggs one to three; or eggs four to nine. The global linear mixed model for the effect of laying order group on egg volume included the following explanatory variables: egg laying date (mean-centred), the total number of eggs laid by each female (mean-centred), and the three-way interaction between laying order group (two-level factor), habitat, and treatment group.

*Nestling body mass on days six and 12 after hatching*

We investigated whether nestling body mass later in the nestling-rearing phase depended on the treatment group and habitat. We created two separate linear mixed models for days six and 12 after hatching, with nestling body mass as the response variable and the following explanatory variables: the number of siblings alive at day two (mean-centred), hatch date (mean-centred), time of day, the two-way interaction between habitat of rearing and treatment group in the nest of rearing, the site of hatching (before cross-fostering was undertaken), and the treatment group in the nest of hatching (before cross-fostering was undertaken). As part of a different study, some hatchlings were cross-fostered and, hence, for the analysis presented here, we included the clutch ID of hatching (a 58-level factor) and the clutch ID of rearing (a 58-level factor) as random effect intercepts in both models on body mass.

*The effect of nestling body mass on nestling survival*

To assess whether nestling body mass two days after hatching influenced the probability of a nestling surviving until fledging, the nestling's survival probability (i.e., the probability that a nestling was alive or dead at each measured time-point; 2-level factor: "fledged" or "dead") was included as a response variable in a binomial generalised linear mixed model. We included the following explanatory variables: hatch date (mean-centred), number of siblings (mean-centred), time of day, and the three-way interaction between the treatment group, nestling body mass two days after hatching, and habitat. The clutch ID of hatching (a 58-level factor) and clutch ID of rearing (a 58-level factor) were included as random effect intercepts.

**Table S1. a)** Likelihood-ratio tests (LRT) for the following predictors explaining the number of eggs produced: the treatment group × habitat interaction and first egg date. Significant P-values are highlighted in bold. "*df*" = degrees of freedom for the LRTs. Superscripts "$^{1}$" and "$^{2}$" refer to linear and quadratic terms, respectively. **b)** Global model coefficients from the linear mixed model of the effects of the following predictors on the number of eggs produced: habitat × treatment group interaction and first egg date. The standard errors and 95% confidence intervals are provided for each model coefficient. N = 61 clutches.

**Number of eggs produced**

**a) Likelihood-ratio test results**

| *Predictors* | $\chi^2$ | *df* | *P-value* |
|---|---|---|---|
| **Habitat** | **14.366** | **1** | **<0.001** |
| **Treatment group** | **7.723** | **1** | **0.005** |
| **$1^{st}$ egg date$^{1}$** | **7.636** | **1** | **<0.001** |
| $1^{st}$ egg date$^{2}$ | 0.126 | 1 | 0.723 |
| Treatment group × Habitat | 2.069 | 1 | 0.150 |

**b) Model coefficients from the global model**

| *Fixed Effects* | *Estimate* | *Standard Error* | *95% CI (Lower* | *Upper)* |
|---|---|---|---|---|
| **Intercept** (Control group in the urban habitat) | 8.188 | 0.458 | **7.271** | **9.105** |
| Treatment group: *Experimental* | 0.517 | 0.632 | -0.749 | 1.783 |
| Habitat: *Forest* | 1.207 | 0.603 | -0.0004 | 2.414 |
| **$1^{st}$ egg laying date$^{1}$** | **-0.147** | **0.047** | **-0.242** | **-0.052** |
| Treatment group: *Experimental* × Habitat: *Forest* | 1.186 | 0.854 | -0.524 | 2.897 |

**Table S2. a)** Likelihood-ratio tests (LRT) for the following predictors explaining the number of eggs produced in the urban habitat: the treatment group and first egg date. Significant P-values are highlighted in bold. "*df*" = degrees of freedom for the LRTs. Superscripts "1" and "2" refer to linear and quadratic terms, respectively. **b)** Model coefficients from the linear mixed model of the effects of the following predictors on the number of eggs produced in the urban habitat: the treatment group and first egg date. The standard errors and 95% confidence intervals are provided for each model coefficient. N = 27 clutches.

| Number of eggs produced in the urban habitat | | | | |
|---|---|---|---|---|
| **a) Likelihood-ratio test results** | | | | |
| ***Predictors*** | $\chi^2$ | ***df*** | ***P-value*** | |
| Treatment group | 0.701 | 1 | 0.403 | |
| 1st egg laying date[1] | 3.108 | 1 | 0.078 | |
| 1st egg laying date[2] | 0.960 | 1 | 0.327 | |
| **b) Model coefficients** | | | | |
| ***Fixed Effects*** | ***Estimate*** | ***Standard Error*** | ***95% CI (Lower \| Upper)*** | |
| **Intercept** (Control group) | 7.941 | 0.426 | **7.061** | **8.821** |
| Treatment group: *Experimental* | 0.472 | 0.594 | -0.754 | 1.697 |
| 1st egg laying date[1] | -0.105 | 0.062 | -0.232 | 0.022 |

**Table S3. a)** Likelihood-ratio tests (LRT) for the following predictors explaining the number of eggs produced in the forest: the treatment group and first egg date. Significant P-values are highlighted in bold. **b)** Model coefficients from the linear mixed model of the effects of the following predictors on the number of eggs produced in the forest: the treatment group and first egg date. The standard errors and 95% confidence intervals are provided for each model coefficient. N = 34 clutches.

| Number of eggs produced in the forest | | | | |
|---|---|---|---|---|
| **a) Likelihood-ratio test results** | | | | |
| ***Predictors*** | $\chi^2$ | ***df*** | ***P-value*** | |
| **Treatment group** | **7.267** | **1** | **0.007** | |
| **1st egg laying date[1]** | **7.110** | **1** | **0.008** | |
| 1st egg laying date[2] | 0.335 | 1 | 0.563 | |
| **b) Minimal model coefficients** | | | | |
| ***Fixed Effects*** | ***Estimate*** | ***Standard Error*** | ***95% CI (Lower \| Upper)*** | |
| **Intercept** (Control group) | 9.643 | 0.408 | 8.812 | 10.476 |
| **Treatment group: *Experimental*** | **1.632** | **0.600** | **0.407** | **2.856** |
| **1st egg laying date[1]** | **-0.192** | **0.062** | **-0.338** | **-0.046** |

**Table S4. a)** Likelihood-ratio tests (LRT) of following predictors explaining egg volume: the treatment group × habitat interaction, number of eggs laid, and first egg date. Clutch ID and egg ID were included as random effects. Significant P-values are highlighted in bold. "*df*" = degrees of freedom for the LRTs. Superscripts "1" and "2" refer to linear and quadratic terms, respectively. **b)** Model coefficients from the Global linear mixed model of the effects of the following predictors on egg volume: habitat × treatment group interaction, the number of eggs laid, and first egg date. The standard errors and 95% confidence intervals (calculated by running 500 parametric bootstrap simulations of the model) are provided for each model coefficient. N = 496 eggs.

| Global model: Egg volume ($mm^3$) | | | | |
|---|---|---|---|---|
| **a) Likelihood-ratio test results** | | | | |
| ***Predictors*** | $\chi^2$ | ***df*** | ***P-value*** | |
| Habitat | - | - | - | |
| Treatment group | - | - | - | |
| Number of eggs laid | 1.995 | 1 | 0.369 | |
| **Egg laying date[1]** | **19.110** | **1** | **<0.001** | |
| Egg laying date[2] | 0.784 | 1 | 0.387 | |
| Habitat × Treatment group | 1.288 | 1 | 0.525 | |
| **b) Model coefficients from the global model** | | | | |
| ***Fixed Effects*** | ***Estimate*** | ***Standard Error*** | ***95% CI (Lower \| Upper)*** | |
| **Intercept** (Control group in the urban habitat) | 1370.036 | 30.663 | **1311.018** | **1430.539** |
| Treatment group: *Experimental* | 25.027 | 37.658 | -55.942 | 102.749 |
| Habitat: *Forest* | -42.002 | 37.434 | -117.973 | 41.654 |
| Number of eggs laid | -7.945 | 7.069 | -20.684 | 5.998 |
| **Egg laying date[1]** | **-5.499** | **1.277** | **-8.222** | **-2.935** |
| Habitat × Treatment group | -36.298 | 49.396 | -126.121 | 62.956 |

| *Random effects* | | |
|---|---|---|
| ***Groups*** | ***Variance*** | ***Standard deviation*** |
| Clutch ID | 5791 | 76.10 |
| Egg ID | 7866 | 88.69 |
| Residual | 1991 | 44.62 |

**Table S5. a)** Likelihood-ratio tests (LRT) of following predictors explaining egg volume: the treatment group, habitat, number of eggs laid, and first egg date. Clutch ID and egg ID were included as random effects. Significant P-values are highlighted in bold. “*df*” = degrees of freedom for the LRTs. Superscripts “1” and “2” refer to linear and quadratic terms, respectively. **b)** Model coefficients from the minimal linear mixed model of the effects of the following predictors on egg volume: habitat, treatment group, the number of eggs laid, and first egg date. The standard errors and 95% confidence intervals (calculated by running 500 parametric bootstrap simulations of the model) are provided for each model coefficient. N=460 eggs.

**Minimal model: Egg volume ($mm^3$)**

**a) Likelihood-ratio test results**

| ***Predictors*** | $\chi^2$ | ***df*** | ***P-value*** |
|---|---|---|---|
| Habitat | 3.745 | 1 | 0.053 |
| Treatment group | 0.029 | 1 | 0.864 |
| Number of eggs laid | 1.638 | 1 | 0.201 |
| **Egg laying date$^{1}$** | **18.304** | **1** | **<0.001** |
| Egg laying date$^{2}$ | 0.784 | 1 | 0.387 |
| Habitat × Treatment group | 1.288 | 1 | 0.525 |

**b) Model coefficients from the minimal model**

| ***Fixed Effects*** | ***Estimate*** | ***Standard Error*** | ***95% CI (Lower \| Upper)*** | |
|---|---|---|---|---|
| **Intercept** (Control group in the urban habitat) | 1379.531 | 27.877 | **1322.991** | **1430.082** |
| Treatment group: *Experimental* | 4.282 | 25.001 | -43.590 | 58.277 |
| Habitat: *Forest* | -58.500 | 29.732 | -119.198 | 4.751 |
| Number of eggs laid | -8.967 | 6.947 | -22.886 | 5.017 |
| **Egg laying date$^{1}$** | **-5.493** | **1.278** | **-8.007** | **-3.179** |

***Random effects***

| ***Groups*** | ***Variance*** | ***Standard deviation*** |
|---|---|---|
| Clutch ID | 7908 | 88.93 |
| Egg ID | 5795 | 76.13 |
| Residual | 1991 | 44.62 |

**Results S1: Total egg volume produced**

The total egg volume produced ($mm^3$) did not depend on the interaction between location and treatment group ($\chi^2_{df=1}$=*1.111, P=0.292; Table S8*). However, this may be an artefact of the small sample size reducing statistical power. When we created within-habitat models, there was no difference in the total volume of eggs produced between the urban treatment groups ($\chi^2_{df=1}$=*0.578, P=0.447; Table S9*). Meanwhile, experimental females produced a greater total volume of eggs than control females in the forest ($\chi^2_{df=1}$=*5.221, P=0.022; Table S10*).

**Table S6. a)** Likelihood-ratio tests (LRT) for the following predictors explaining the total egg volume produced: the treatment group × habitat interaction and first egg date. Significant P-values are highlighted in bold and italics. "*df*" = degrees of freedom for the LRTs. Superscripts "1" and "2" refer to linear and quadratic terms, respectively. **b)** Global model coefficients from the global linear model of the effects of the following predictors on the total volume of egg material produced: the habitat × treatment group interaction and first egg date. The standard errors and 95% confidence intervals are provided for each model coefficient N = 61 clutches.

| Global model: Total volume of egg ($mm^3$) produced by a female | | | | |
|---|---|---|---|---|
| **a) Likelihood-ratio test results** | | | | |
| ***Predictors*** | $\chi^2$ | ***df*** | ***P-value*** | |
| Habitat | - | - | - | |
| Treatment group | - | - | - | |
| **1st egg laying date[1]** | **11.815** | **1** | **<0.001** | |
| 1st egg laying date[2] | 0.377 | 1 | 0.539 | |
| Treatment group × Habitat | 1.111 | 1 | 0.292 | |
| **b) Model coefficients from the global model** | | | | |
| ***Fixed Effects*** | ***Estimate*** | ***Standard Error*** | ***95% CI (Lower \| Upper)*** | |
| **Intercept** (Control group in the urban habitat) | 11299.800 | 664.200 | 9967.674 | 12631.962 |
| Treatment group: *Experimental* | 678.980 | 919.690 | -398.582 | 3095.857 |
| Habitat: *Forest* | 1348.685 | 871.180 | -1169.499 | 2527.049 |
| **1st egg laying date** | **-235.318** | **68.130** | **-371.680** | **-98.912** |
| Treatment group: *Experimental* × Habitat: *Forest* | 1241.930 | 1226.250 | -1217.545 | 3701.334 |

**Table S7. a)** Likelihood-ratio tests (LRT) for the following predictors explaining the total volume of egg produced by a female in the urban habitat: treatment group and first egg date. Significant P-values are highlighted in bold. "*df*" = degrees of freedom for the LRTs. Superscripts "1" and "2" refer to linear and quadratic terms, respectively. **b)** Model coefficients from the linear model of the effects of the following predictors on total volume of egg produced in the urban habitat: treatment group and first egg date. N = 27 clutches.

| Total volume of egg ($mm^3$) produced by a female in the urban habitat | | | | |
|---|---|---|---|---|
| **a) Likelihood-ratio test results** | | | | |
| ***Predictors*** | $\chi^2$ | ***df*** | ***P-value*** | |
| Treatment group | 0.578 | 1 | 0.447 | |
| **1st egg laying date[1]** | **5.352** | **1** | **0.021** | |
| 1st egg laying date[2] | 0.077 | 1 | 0.078 | |
| **b) Model coefficients** | | | | |
| ***Fixed Effects*** | ***Estimate*** | ***Standard Error*** | ***95% CI (Lower \| Upper)*** | |
| **Intercept** (Control group in the urban habitat) | 10889.770 | 646.201 | 9549.624 | 12229.913 |
| Treatment group: *Experimental* | 645.830 | 900.303 | -1221.291 | 2512.942 |
| **1st egg date[1]** | **-210.932** | **92.056** | **-401.827** | **-20.041** |

**Table S8. a)** Likelihood-ratio tests (LRT) for the the following predictors explaining the total volume of egg produced by a female in the forest: treatment group and first egg date. Significant P-values are highlighted in bold. "*df*" = degrees of freedom for the LRTs. Superscripts "1" and "2" refer to linear and quadratic terms, respectively. **b)** Model coefficients from the linear model of the effects of the following predictors on total volume of egg material produced in the forest: treatment group, and first egg date. N = 34 clutches.

| Total volume of egg ($mm^3$) produced by a female in the forest | | | | |
|---|---|---|---|---|
| **a) Likelihood-ratio test results** | | | | |
| ***Predictors*** | $\chi^2$ | ***df*** | ***P-value*** | |
| **Treatment group** | **5.221** | **1** | **0.022** | |
| **1st egg date[1]** | **6.640** | **1** | **0.010** | |
| 1st egg date[2] | 0.839 | 1 | 0.360 | |
| **b) Model coefficients** | | | | |
| ***Fixed Effects*** | ***Estimate*** | ***Standard Error*** | ***95% CI (Lower \| Upper)*** | |
| **Intercept** (Control group in the forest) | 12988.400 | 575.810 | 11812.436 | 14168.036 |
| **Treatment group: *Experimental*** | **1887.202** | **832.208** | **187.661** | **3586.797** |
| **1st egg date[1]** | **-261.520** | **101.100** | **-467.997** | **-54.964** |

**Table S9. a)** Likelihood-ratio tests (LRT) for the minimal model of the following predictors explaining the total volume of egg produced by a female: treatment group, habitat, and first egg date. Significant P-values are highlighted in bold. "*df*" = degrees of freedom for the LRTs. Superscripts "1" and "2" refer to linear and quadratic terms, respectively. **b)** Minimal model coefficients from the linear model of the effects of the following predictors on total volume of egg material produced: habitat, treatment group, and first egg date. The standard errors and 95% confidence intervals are provided for each model coefficient. N=61 clutches.

| Minimal model: Total volume of egg ($mm^3$) produced by a female | | | | |
|---|---|---|---|---|
| **a) Likelihood-ratio test results** | | | | |
| ***Predictors*** | $\chi^2$ | ***df*** | ***P-value*** | |
| **Habitat** | **9.049** | **1** | **0.003** | |
| **Treatment group** | **5.437** | **1** | **0.020** | |
| **1st egg date¹** | **12.773** | **1** | **<0.001** | |
| 1st egg date² | 0.392 | 1 | 0.531 | |
| Treatment group × Habitat | 1.112 | 1 | 0.292 | |
| **b) Model coefficients** | | | | |
| ***Fixed Effects*** | ***Estimate*** | ***Standard Error*** | ***95% CI (Lower \| Upper)*** | |
| **Intercept** (Control group in the urban habitat) | 10951.36 | 568.240 | 9812.110 | 12090.6121 |
| **Treatment group:** ***Experimental*** | **1943.360** | **600.880** | **179.584** | **2588.966** |
| **Habitat:** ***Forest*** | **1943.36** | **643.59** | **653.030** | **2588.966** |
| **1st egg date¹** | **-245.390** | **67.280** | **-380.272** | **-110.499** |

#### *Results S2: The effect of laying order, treatment group, and habitat on egg volume*

When eggs were grouped according to their position in the laying sequence (1. eggs one to three or 2. four to nine), there was no three-way interaction between the laying order group, treatment group, and habitat ($\chi^2_{df=1}=3.119$, $P=0.077$). However, we found that egg volume depended on the two-way interactions between location and laying order ($\chi^2_{df=1}=10.499$, $P=0.001$) and treatment group and laying order ($\chi^2_{df=1}=4.394$, $P=0.036$). There was no difference in volume between habitats for eggs one to three in the lay sequence. However, eggs four to nine in the forest were smaller than those in the urban habitat. Additionally, eggs one to three were larger in experimental nests than control nests. Egg volume for eggs four to nine in the lay sequence did not differ between treatment groups.

**Table S10. a)** Likelihood-ratio tests (LRT) for the following predictors explaining egg volume: the treatment group × habitat × laying order group interaction, number of eggs laid, and first egg date. Clutch ID and egg ID were included as random effects. Significant P-values are highlighted in bold. “*df*” = degrees of freedom for the LRTs. Superscripts “$^1$” and “$^2$” refer to linear and quadratic terms, respectively. **b)** Global model coefficients from the global linear mixed model of the effects of the following predictors on egg volume: treatment group × habitat × laying order group interaction, the number of eggs laid, and first egg date. The standard errors and 95% confidence intervals (calculated by running 500 parametric bootstrap simulations of the model) are provided for each model coefficient. N = 460 eggs.

| Global model: Egg volume ($mm^3$) | | | | |
|---|---|---|---|---|
| **a) Likelihood-ratio test results** | | | | |
| ***Predictors*** | $\chi^2$ | ***df*** | | ***P-value*** |
| Habitat | - | - | | - |
| Treatment group | - | - | | - |
| Lay order group | - | - | | - |
| Number of eggs laid | 1.677 | 1 | | 0.195 |
| **Egg laying date$^1$** | **12.248** | **1** | | **<0.001** |
| Egg laying date$^2$ | 0.609 | 1 | | 0.435 |
| Habitat × lay order group × Treatment group | 3.119 | 1 | | 0.077 |
| **b) Model coefficients from the global model** | | | | |
| ***Fixed Effects*** | ***Estimate*** | ***Standard Error*** | ***95% CI (Lower \| Upper)*** | |
| **Intercept** (Eggs 1-3 from the control group in the urban habitat) | 1339.870 | 35.894 | 1266.660 | 1424.988 |
| Treatment group: *Experimental* | 33.851 | 44.145 | -51.495 | 116.961 |
| Habitat: *Forest* | -23.908 | 43.705 | -114.369 | 62.364 |
| Number of eggs laid | -9.672 | 7.423 | -24.009 | 4.380 |
| **Egg laying date$^1$** | **-6.574** | **1.872** | **-10.795** | **-3.024** |
| Lay order group: *Eggs 4-9* | 41.044 | 24.624 | -8.034 | 89.311 |
| Habitat: *Forest* × Lay order group: *Eggs 4-9* | -26.862 | 28.870 | -86.928 | 25.314 |
| Lay order group: *Eggs 4-9* × Treatment group: *Experimental* | -0.989 | 29.682 | -55.545 | 57.222 |
| Habitat: *Forest* × Treatment group: *Experimental* | 23.267 | 58.816 | -87.518 | 124.313 |
| Habitat: *Forest* × Lay order group: *Eggs 4-9* × Treatment group: *Experimental* | -69.833 | 39.354 | -147.739 | 11.559 |

| ***Random effects*** | | |
|---|---|---|
| ***Groups*** | ***Variance*** | ***Standard deviation*** |
| Clutch ID | 8396 | 1.63 |
| Egg ID | 5472 | 73.97 |
| Residual | 2048 | 45.26 |

**Table S11. a)** Likelihood-ratio tests (LRT) from the minimal model for the following predictors explaining egg volume: the treatment group × habitat interaction, the treatment group × laying order group interaction, the habitat × laying order group interaction, the number of eggs laid, and egg laying date. Clutch ID and egg ID were included as random effects. Significant P-values are highlighted in bold. "*df*" = degrees of freedom for the LRTs. Superscripts "1" and "2" refer to linear and quadratic terms, respectively. **b)** Minimal model coefficients from the linear mixed model of the effects of the following predictors on egg volume: Treatment group × laying order interaction, Habitat × laying order interaction, experimental group, the number of eggs laid, and first egg date. The standard errors and 95% confidence intervals are provided for each model coefficient. N=460 eggs.

| Minimal model: Egg volume ($mm^3$) | | | |
|---|---|---|---|
| **a) Likelihood-ratio test results** | | | |
| ***Predictors*** | $\chi^2$ | ***df*** | ***P-value*** |
| Habitat | - | - | - |
| Treatment group | - | - | - |
| Lay order group | - | - | - |
| Number of eggs laid | 2.088 | 1 | 0.149 |
| **Egg laying date$^1$** | **12.665** | **1** | **<0.001** |
| Egg laying date$^2$ | 1.111 | 1 | 0.292 |
| **Habitat × Lay order group** | **10.499** | **1** | **0.001** |
| **Lay order group × Treatment group** | **4.394** | **1** | **0.036** |
| Habitat × Treatment group | 0.322 | 1 | 0.571 |
| Habitat × lay order group × Treatment group | 3.119 | 1 | 0.077 |

| **b) Model coefficients from the minimal model** | | | | |
|---|---|---|---|---|
| ***Fixed Effects*** | ***Estimate*** | ***Standard Error*** | ***95% CI (Lower \| Upper)*** | |
| **Intercept** (Eggs 1-3 from the control group in the urban habitat) | 1328.949 | 31.356 | 1267.540 | 1387.274 |
| Treatment group: *Experimental* | 46.108 | 29.566 | -13.501 | 104.300 |
| Habitat: *Forest* | -9.110 | 33.608 | -74.997 | 52.123 |
| Number of eggs laid | -10.574 | 7.265 | -25.910 | 2.952 |
| **Egg laying date$^1$** | **-6.682** | **1.872** | **-10.495** | **-2.813** |
| **Lay order group: *Eggs 4-9*** | **66.061** | **20.263** | **27.719** | **112.774** |
| **Habitat: *Forest* × Lay order group: *Eggs 4-9*** | **-64.453** | **19.741** | **-99.789** | **-26.054** |
| **Treatment group: *Experimental* × Lay order group: *Eggs 4-9*** | **-41.164** | **19.580** | **-84.300** | **-4.554** |

| ***Random effects*** | | |
|---|---|---|
| ***Groups*** | ***Variance*** | ***Standard deviation*** |
| Clutch ID | 8283 | 91.01 |
| Egg ID | 5552 | 74.51 |
| Residual | 2048 | 45.25 |

**Table S12. a)** Likelihood-ratio tests (LRT) for the following predictors explaining egg volume: the treatment group × habitat × laying order (fitted linearly) interaction, number of eggs laid, and first egg date. Clutch ID and egg ID were included as random effects. Significant P-values are highlighted in bold. "*df*" = degrees of freedom for the LRTs. Superscripts "1" and "2" refer to linear and quadratic terms, respectively. **b)** Global model coefficients and confidence intervals from the linear mixed model of the effects of the following predictors on egg volume: treatment group × habitat × laying order interaction, the number of eggs laid, and first egg date. The standard errors and 95% confidence intervals (calculated by running 500 parametric bootstrap simulations of the model) are provided for each model coefficient. N=460 eggs.

| Egg volume ($mm^3$) | | | | |
|---|---|---|---|---|
| **a) Likelihood-ratio test results** | | | | |
| ***Predictors*** | $x^2$ | ***df*** | | ***P-value*** |
| Habitat | - | - | | - |
| Treatment group | - | - | | - |
| Lay order | - | - | | - |
| Number eggs laid | 2.916 | 1 | | 0.088 |
| **Egg laying date$^1$** | **13.183** | **1** | | **<0.001** |
| Egg laying date$^2$ | 0.860 | 1 | | 0.354 |
| **Habitat × lay order × Treatment group** | **3.949** | **1** | | **0.047** |
| **b) Model coefficients** | | | | |
| ***Fixed Effects*** | ***Estimate*** | ***Standard Error*** | ***95% CI (Lower*** | ***Upper)*** |
| **Intercept** (1st laid egg in the control group in the urban habitat) | 1369.831 | 31.632 | 1306.625 | 1432.789 |
| Treatment group: *Experimental* | 37.509 | 40.022 | -41.787 | 117.920 |
| Habitat: *Forest* | -45.204 | 39.334 | -123.547 | 33.309 |
| Number of eggs laid | -13.580 | 7.928 | -29.636 | 2.036 |
| **Egg laying date$^1$** | **-9.599** | **2.630** | **-15.140** | **-4.377** |
| Lay order | 11.619 | 5.697 | 0.391 | 22.961 |
| Habitat: *Forest* × Lay order | -3.521 | 5.851 | -10.354 | 7.969 |
| Lay order × Treatment group: *Experimental* | 1.796 | 6.171 | -10.354 | 13.918 |
| Habitat: *Forest* × Treatment group: *Experimental* | -29.393 | 52.312 | -134.151 | 74.156 |
| **Habitat: *Forest* × Lay order × Treatment group: *Experimental*** | **-15.443** | **7.729** | **-30.626** | **-0.213** |

| ***Random effects*** | | |
|---|---|---|
| ***Groups*** | ***Variance*** | ***Standard deviation*** |
| Clutch ID | 8961 | 94.66 |
| Egg ID | 5415 | 73.58 |
| Residual | 2039 | 45.15 |

**Table S13. a)** Likelihood-ratio tests (LRT) for the following predictors explaining nestling body mass two days after hatching: the treatment group × habitat interaction, number of siblings in the nest, time of day, and hatch date. Clutch ID was included as a random effect. Significant P-values are highlighted in bold. "*df*" = degrees of freedom for the LRTs. Superscripts "1" and "2" refer to linear and quadratic terms, respectively. **b)** Global model coefficients from the linear mixed model of the effects of the following predictors on nestling mass two days after hatching: treatment group × habitat interaction, number of siblings in the nest, time of day, and hatch date. The standard errors and 95% confidence intervals (calculated by running 500 parametric bootstrap simulations of the model) are provided for each model coefficient. N = 372 nestlings.

| Global model: Nestling mass (g) two days after hatching | | | | |
|---|---|---|---|---|
| **a) Likelihood-ratio test results** | | | | |
| ***Predictors*** | $\chi^2$ | ***df*** | | ***P-value*** |
| Habitat | - | - | | - |
| Treatment group | - | - | | - |
| Hatch date[1] | 0.189 | 1 | | 0.664 |
| Hatch date[2] | 0.219 | 1 | | 0.640 |
| Number of siblings | 0.725 | 1 | | 0.395 |
| Time of day | 1.757 | 1 | | 0.185 |
| Treatment group × Habitat | 0.132 | 1 | | 0.717 |
| **b) Model coefficients from the global model** | | | | |
| ***Fixed Effects*** | ***Estimate*** | ***Standard Error*** | ***95% CI (Lower*** | ***Upper)*** |
| **Intercept** (Control group in the urban habitat in the morning) | 1.928 | 0.100 | 1.720 | 2.084 |
| Treatment group: *Experimental* | -0.294 | 0.167 | -0.608 | 0.047 |
| Habitat: *Forest* | 0.123 | 0.141 | -0.118 | 0.361 |
| Number of siblings | -0.024 | 0.028 | -0.077 | 0.008 |
| Hatch date[1] | 0.006 | 0.014 | -0.022 | 0.030 |
| Time of day: *Afternoon* | 0.141 | 0.104 | -0.039 | 0.310 |
| Treatment group: *Experimental* × Habitat: *Forest* | 0.071 | 0.196 | -0.295 | 0.414 |

| ***Random effects*** | | |
|---|---|---|
| ***Groups*** | ***Variance*** | ***Standard deviation*** |
| Clutch ID | 0.097 | 0.312 |
| Residual | 0.156 | 0.395 |

**Table S14. a)** Likelihood-ratio tests (LRT) for the following predictors explaining nestling body mass two days after hatching: treatment group, habitat, number of siblings in the nest, time of day, and hatch date. Clutch ID was included as a random effect. Significant P-values are highlighted in bold. "*df*" = degrees of freedom for the LRTs. Superscripts "1" and "2" refer to linear and quadratic terms, respectively. **b)** Minimal model coefficients from the linear mixed model of the effects of the following predictors on nestling mass two days after hatching: treatment group, habitat, number of siblings in the nest, time of day, and hatch date. The standard errors and 95% confidence intervals (calculated by running 500 parametric bootstrap simulations of the model) are provided for each model coefficient. N=372 nestlings.

| Minimal Model: Nestling mass (g) two days after hatching | | | |
|---|---|---|---|
| **a) Likelihood-ratio test results** | | | |
| ***Predictors*** | $x^2$ | ***df*** | ***P-value*** |
| Habitat | 1.730 | 1 | 0.188 |
| **Treatment group** | **4.117** | **1** | **0.042** |
| Hatch date$^1$ | 0.182 | 1 | 0.670 |
| Hatch date$^2$ | 0.219 | 1 | 0.640 |
| Number of siblings | 0.681 | 1 | 0.409 |
| Time of day | 1.803 | 1 | 0.179 |
| Treatment group × Habitat | 0.132 | 1 | 0.717 |

| **b) Model coefficients from the minimal model** | | | | |
|---|---|---|---|---|
| ***Fixed Effects*** | ***Estimate*** | ***Standard Error*** | ***95% CI (Lower*** | ***Upper)*** |
| **Intercept** (Control group in the urban habitat in the morning) | 1.912 | 0.091 | 1.749 | 2.059 |
| **Treatment group: *Experimental*** | **-0.252** | **0.121** | **-0.479** | **-0.045** |
| Habitat: *Forest* | 0.153 | 0.115 | -0.057 | 0.324 |
| Number of siblings | -0.023 | 0.028 | -0.086 | 0.025 |
| Hatch date | 0.006 | 0.014 | -0.021 | 0.028 |
| Time of day: *Afternoon* | 0.143 | 0.104 | -0.086 | 0.318 |

| ***Random effects*** | | |
|---|---|---|
| ***Groups*** | ***Variance*** | ***Standard deviation*** |
| Clutch ID | 0.098 | 0.312 |
| Residual | 0.156 | 0.395 |

***Results S3: The effect of treatment group on nestling body mass on days six and 12 after hatching***

Only on day 12 after hatching was there was an interaction between treatment group and habitat on nestling body mass, with there being no interaction effect on day six (day six after hatching: $\chi^2_{df=1}=1.342$, $P=0.247$; Table S15-S16, day 12 after hatching: $\chi^2_{df=1}=1.746$, $P<0.001$, Table S17). In the forest, there were no differences in nestling body mass between treatment groups on days six and 12, the same holds for both treatment groups in the urban habitat on day six. Meanwhile, in the urban habitat, nestlings in experimental nests were heavier than those in control nests 12 days after hatching.

**Table S15. a)** Likelihood-ratio tests (LRT) from the global model for the following predictors explaining nestling body mass six days after hatching: the treatment group in the nest of rearing × the habitat of the nest of rearing interaction, treatment group in the nest of hatching, habitat in the nest of hatching, the number of siblings in the nest, time of day, and hatch date. Clutch ID of rearing and Clutch ID of hatching were included as random effects. Significant P-values are highlighted in bold. "*df*" = degrees of freedom for the LRTs. Superscripts "1" and "2" refer to linear and quadratic terms, respectively. **b)** Global model coefficients from the linear mixed model of the effects of the following predictors on nestling mass six days after hatching: treatment group in the nest of rearing× habitat in the nest of rearing interaction, the treatment group in the nest of hatching, habitat in the nest of hatching, number of siblings in the nest, time of day, and hatch date. The standard errors and 95% confidence intervals (calculated by running 500 parametric bootstrap simulations of the model) are provided for each model coefficient. N=353 nestlings.

| Global model: Nestling mass (g) six days after hatching | | | | |
|---|---|---|---|---|
| **a) Likelihood-ratio test results** | | | | |
| ***Predictors*** | $\chi^2$ | ***df*** | ***P-value*** | |
| Treatment group in the nest of hatching | 0.415 | 1 | 0.519 | |
| Treatment group in the nest of rearing | - | - | - | |
| Habitat in the nest of hatching | 0.400 | 1 | 0.527 | |
| Habitat in the nest of rearing | - | - | - | |
| Hatch date$^1$ | 0.778 | 1 | 0.378 | |
| Hatch date$^2$ | 0.219 | 1 | 0.640 | |
| Number of siblings | 0.430 | 1 | 0.512 | |
| Time of day | 0.958 | 1 | 0.328 | |
| Treatment group in the nest of rearing × Habitat in the nest of rearing | 1.342 | 1 | 0.247 | |
| **b) Global Model coefficients** | | | | |
| ***Fixed Effects*** | ***Estimate*** | ***Standard Error*** | ***95% CI (Lower*** | ***Upper)*** |
| **Intercept** (Control group in the urban habitat in the morning) | 4.688 | 0.271 | 4.128 | 5.116 |
| Treatment group in the nest of hatching: *Experimental* | -0.210 | 0.325 | -0.949 | 0.358 |
| Treatment group in the nest of rearing: *Experimental* | 0.645 | 0.470 | -0.378 | 1.417 |
| Habitat in the nest of hatching: *Forest* | 0.146 | 0.226 | -0.266 | 0.533 |
| **Habitat in the nest of rearing: *Forest*** | **1.041** | **0.323** | **0.428** | **1.606** |
| Number of siblings | -0.023 | 0.028 | -0.174 | 0.054 |
| Hatch date | -0.042 | 0.063 | -0.103 | 0.033 |
| Time of day: *Afternoon* | 0.235 | 0.239 | -0.208 | 0.654 |
| Treatment group in the nest of rearing: *Experimental*× Habitat in the nest of rearing: *Forest* | -0.536 | 0.459 | -1.450 | 0.247 |

| ***Random effects*** | | |
|---|---|---|
| ***Groups*** | ***Variance*** | ***Standard deviation*** |
| Clutch ID of rearing | 0.404 | 0.636 |
| Clutch ID of hatching | 0.342 | 0.585 |
| Residual | 0.637 | 0.798 |

**Table S16. a)** Likelihood-ratio tests (LRT) from the global model for the following predictors explaining nestling body mass six days after hatching: the treatment group in the nest of rearing × the habitat of the nest of rearing interaction, treatment group in the nest of hatching, habitat in the nest of hatching, the number of siblings in the nest, time of day, and hatch date. Clutch ID of rearing and Clutch ID of hatching were included as random effects. Significant P-values are highlighted in bold. "*df*" = degrees of freedom for the LRTs. Superscripts "1" and "2" refer to linear and quadratic terms, respectively. **b)** Minimal model coefficients from the linear mixed model of the effects of the following predictors on nestling mass six days after hatching: treatment group in the nest of rearing, habitat in the nest of rearing, the treatment group in the nest of hatching, habitat in the nest of hatching, number of siblings in the nest, time of day, and hatch date. The standard errors and 95% confidence intervals (calculated by running 500 parametric bootstrap simulations of the model) are provided for each model coefficient. N=353 nestlings.

| Nestling mass (g) six days after hatching | | | | |
|---|---|---|---|---|
| **a) Likelihood-ratio test results** | | | | |
| ***Predictors*** | $\chi^2$ | ***df*** | | ***P-value*** |
| Treatment group in the nest of hatching | 0.379 | 1 | | 0.538 |
| Treatment group in the nest of rearing | 0.655 | 1 | | 0.418 |
| Habitat in the nest of hatching | 0.375 | 1 | | 0.541 |
| **Habitat in the nest of rearing** | **8.097** | **1** | | **0.004** |
| Hatch date$^1$ | 0.810 | 1 | | 0.368 |
| Hatch date$^2$ | 0.219 | 1 | | 0.640 |
| Number of siblings | 0.547 | 1 | | 0.460 |
| Time of day | 0.743 | 1 | | 0.389 |
| Treatment group in the nest of rearing × Habitat in the nest of rearing | 1.342 | 1 | | 0.247 |
| **b) Model coefficients from the minimal model** | | | | |
| ***Fixed Effects*** | ***Estimate*** | ***Standard Error*** | ***95% CI (Lower*** | ***Upper)*** |
| **Intercept** (Control group in the urban habitat in the morning) | 4.819 | 0.250 | 4.282 | 5.220 |
| Treatment group in the nest of hatching: *Experimental* | -0.201 | 0.373 | -0.846 | 0.329 |
| Treatment group in the nest of rearing: *Experimental* | 0.303 | 0.470 | -0.421 | 0.950 |
| Habitat in the nest of hatching: *Forest* | 0.141 | 0.226 | -0.236 | 0.512 |
| **Habitat in the nest of rearing: *Forest*** | **0.835** | **0.276** | **0.254** | **1.286** |
| Number of siblings | -0.048 | 0.064 | -0.169 | 0.066 |
| Hatch date | -0.033 | 0.036 | -0.103 | 0.033 |
| Time of day: *Afternoon* | 0.209 | 0.241 | -0.099 | 0.611 |

| ***Random effects*** | | |
|---|---|---|
| ***Groups*** | ***Variance*** | ***Standard deviation*** |
| Clutch ID of rearing | 0.423 | 0.650 |
| Clutch ID of hatching | 0.341 | 0.584 |
| Residual | 0.637 | 0.798 |

**Table S17. a)** Likelihood-ratio tests (LRT) from the global model for the following predictors explaining nestling body mass 12 days after hatching: the treatment group in the nest of rearing × the habitat of the nest of rearing interaction, treatment group in the nest of hatching, habitat in the nest of hatching, the number of siblings in the nest, time of day, and hatch date. Clutch ID of rearing and Clutch ID of hatching were included as random effects. Significant P-values are highlighted in bold. "*df*" = degrees of freedom for the LRTs. Superscripts "$^{1}$" and "$^{2}$" refer to linear and quadratic terms, respectively. **b)** Global model coefficients from the linear mixed model of the effects of the following predictors on nestling mass 12 days after hatching: treatment group in the nest of rearing × habitat in the nest of rearing interaction, the treatment group in the nest of hatching, habitat in the nest of hatching, number of siblings in the nest, time of day, and hatch date. The standard errors and 95% confidence intervals (calculated by running 500 parametric bootstrap simulations of the model) are provided for each model coefficient. N=353 nestlings.

**Nestling mass (g) 12 days after hatching**

**a) Likelihood-ratio test results**

| *Predictors* | $\chi^2$ | *df* | *P-value* |
|---|---|---|---|
| Treatment group in the nest of hatching | 1.111 | 1 | 0.292 |
| **Treatment group in the nest of rearing** | - | - | - |
| Habitat in the nest of hatching | 1.042 | 1 | 0.308 |
| **Habitat in the nest of rearing** | **-** | **-** | **-** |
| **Hatch date$^{1}$** | **6.320** | **1** | **0.012** |
| Hatch date$^{2}$ | 0.384 | 1 | 0.536 |
| Number of siblings | 0.501 | 1 | 0.479 |
| Time of day | 2.194 | 1 | 0.139 |
| **Treatment group in the nest of rearing × Habitat in the nest of rearing** | **1.746** | **1** | **<0.001** |

**b) Model coefficients from the minimal model**

| *Fixed Effects* | *Estimate* | *Standard Error* | *95% CI (Lower* | *Upper)* |
|---|---|---|---|---|
| **Intercept** (Control group in the urban habitat in the morning) | 7.814 | 0.347 | 7.138 | 8.356 |
| Treatment group in the nest of hatching: *Experimental* | -0.290 | 0.274 | -0.864 | 0.131 |
| **Treatment group in the nest of rearing: *Experimental*** | **2.225** | **0.542** | **1.110** | **3.132** |
| **Habitat in the nest of hatching: *Forest*** | **-0.198** | **0.193** | **1.906** | **0.111** |
| Habitat in the nest of rearing: *Forest* | 2.656 | 0.420 | 0.254 | 3.338 |
| Number of siblings | 0.050 | 0.070 | -0.102 | 0.154 |
| **Hatch date** | **-0.087** | **0.034** | **-0.153** | **-0.030** |
| Time of day: *Afternoon* | 0.393 | 0.260 | -0.145 | 0.836 |
| **Treatment group in the nest of rearing: *Experimental* × Habitat in the nest of rearing: *Forest*** | **-1.593** | **0.534** | **-2.644** | **-0.702** |

***Random effects***

| *Groups* | *Variance* | *Standard deviation* |
|---|---|---|
| Clutch ID of rearing | 0.423 | 0.650 |
| Clutch ID of hatching | 0.189 | 0.435 |
| Residual | 0.438 | 0.662 |

#### *Results S4: The effect of nestling body mass two days after hatching on survival*

We found no differential effect of nestling mass on survival two days after hatching between treatment groups and habitats, as represented by the absence of a three-way interaction between nestling body mass two days after hatching, treatment group, and habitat ($\chi^2_{df=1}=0.256$, *P=0.611;* Table S18). In both treatment groups and habitats, the likelihood of a nestling surviving until fledging increased with body mass two days after hatching ($\chi^2_{df=1}=10.647$, *P=0.001*; Table S19).

**Table S18. a)** Likelihood-ratio tests (LRT) for the following predictors explaining the nestlings fate: the nestling body mass two days after hatching × treatment group × habitat interaction, number of siblings, time of day, and hatch date. N= 364. Clutch of rearing and hatching were included as random effects. Significant P-values are highlighted in bold. "*df*" = degrees of freedom for the LRTs. Superscripts "1" and "2" refer to linear and quadratic terms, respectively. **b)** Global model coefficients from the binomial generalised linear mixed model of the effects of the following predictors on the nestling's fate: the interaction between nestling body mass two days after hatching × the treatment group × habitat, number of siblings, time of day, and hatch date. The standard errors and 95% confidence intervals are provided for each model coefficient. N= 364 nestlings.

**Global model: Fate (Dead or Fledged)**

a) **Likelihood-ratio test results**

| ***Predictors*** | $\chi^2$ | ***df*** | ***P-value*** |
|---|---|---|---|
| Habitat | - | - | - |
| Treatment group | - | - | - |
| Mass (g) two days after hatching | - | - | - |
| Number of siblings | 2.142 | 1 | 0.143 |
| **Hatch date[1]** | **4.295** | **1** | **0.038** |
| Hatch date[2] | 2.296 | 1 | 0.130 |
| Time of day | 3.545 | 1 | 0.060 |
| Mass two days after hatching × treatment group × Habitat | 0.256 | 1 | 0.611 |

b) **Global Model coefficients**

| ***Fixed Effects*** | ***Estimate*** | ***Standard Error*** | ***95% CI (Lower*** | ***Upper)*** |
|---|---|---|---|---|
| **Intercept** (Control group in the urban habitat) | -4.123 | 0.573 | -9.538 | 1.291 |
| Mass (g) two days after hatching | 1.087 | 0.993 | -0.859 | 3.032 |
| Treatment group: *Experimental* | 3.003 | 3.960 | -4.759 | 10.764 |
| Habitat: *Forest* | 5.555 | 4.016 | -2.317 | 13.426 |
| **Hatch date[1]** | **-0.531** | **0.271** | **-1.062** | **-0.001** |
| Number of siblings | 0.691 | 0.464 | -0.219 | 1.600 |
| Time of day: *Afternoon* | -3.424 | 1.841 | -7.032 | 0.184 |
| Mass two days after hatching × Habitat: *Forest* | 1.355 | 1.523 | -1.630 | 4.341 |
| Mass two days after hatching × Treatment group: *Experimental* | 0.691 | 1.475 | -2.201 | 3.582 |
| Treatment group: *Experimental* × Habitat: *Forest* | -3.829 | 6.093 | -15.772 | 8.112 |
| Mass two days after hatching × Habitat: *Forest* × Treatment group: *Experimental* | 1.515 | 3.038 | -4.439 | 7.469 |

***Random effects***

| ***Groups*** | ***Variance*** | ***Standard deviation*** |
|---|---|---|
| Clutch ID of hatching | 5.469 | 2.339 |
| Clutch ID of rearing | 21.318 | 4.617 |

**Table S19. a)** Likelihood-ratio tests (LRT) from the minimal model for the following predictors explaining the nestlings fate: nestling body mass two days after hatching, the treatment group, habitat, number of siblings, time of day, and hatch date. Clutch of hatching and rearing were included as random effects. Significant P-values are highlighted in bold. “*df*” = degrees of freedom for the LRTs. Superscripts “$^{1}$” and “$^{2}$” refer to linear and quadratic terms, respectively. **b)** Model coefficients from the minimal binomial generalised linear mixed model of the effects of the following predictors on the nestling’s fate: nestling body mass two days after hatching, the treatment group, habitat, number of siblings, time of day, and hatch date. The standard errors and 95% confidence intervals are provided for each model coefficient. N=364 nestlings.

**Minimal model: Fate (Dead or Fledged)**

**a) Likelihood-ratio test results**

| *Predictors* | $x^2$ | *df* | *P-value* |
|---|---|---|---|
| **Mass (g) two days after hatching** | **10.647** | **1** | **0.001** |
| **Habitat** | **17.562** | **1** | **<0.001** |
| Treatment group | 2.542 | 1 | 0.111 |
| Number of siblings | 1.708 | 1 | 0.191 |
| **Hatch date$^{1}$** | **4.222** | **1** | **0.039** |
| Hatch date$^{2}$ | 2.296 | 1 | 0.130 |
| Time of day | 3.537 | 1 | 0.060 |
| Mass (g) at day two × Habitat | 1.631 | 1 | 0.202 |
| Habitat × treatment group | 0.147 | 1 | 0.702 |
| Mass (g) at day two × treatment group | 3.233 | 1 | 0.199 |
| Mass (g) at day two × treatment group × Habitat | 0.256 | 1 | 0.611 |

**b) Model coefficients from the minimal model**

| *Fixed Effects* | *Estimate* | *Standard Error* | *95% CI (Lower* | *Upper)* |
|---|---|---|---|---|
| **Intercept** (Control group in the urban habitat) | -5.653 | 2.260 | -10.082 | -1.223 |
| **Mass (g) two days after hatching** | **1.987** | **0.689** | **0.637** | **3.337** |
| Treatment group: *Experimental* | 3.196 | 2.005 | -0.734 | 7.126 |
| **Habitat: *Forest*** | **7.660** | **2.491** | **2.777** | **12.543** |
| **Hatch date$^{1}$** | **-0.522** | **0.262** | **-1.036** | **-0.007** |
| Number of siblings | 0.588 | 0.435 | -0.219 | 1.600 |
| Time of day: *Afternoon* | -3.285 | 1.762 | -6.737 | 0.168 |

***Random effects***

| *Groups* | *Variance* | *Standard deviation* |
|---|---|---|
| Clutch of hatching | 5.764 | 2.401 |
| Clutch of rearing | 21.369 | 4.623 |

**Table S20. a)** Likelihood-ratio tests (LRT) for the following predictors explaining the number of nestlings alive: the treatment group × habitat × age interaction, clutch size, and hatch date. Clutch ID was included as a random effect. Significant P-values are highlighted in bold. "*df*" = degrees of freedom for the LRTs. Superscripts "1" and "2" refer to linear and quadratic terms, respectively. **b)** Global model coefficients from the linear mixed model of the effects of the following predictors on the number of chicks alive: the treatment group × habitat × age interaction, clutch size, and hatch date. The standard errors and 95% confidence intervals (calculated by running 500 parametric bootstrap simulations of the model) are provided for each model coefficient. N=60 broods.

**Number of nestlings alive**

### a) Likelihood-ratio test results

| *Predictors* | $\chi^2$ | *df* | *P-value* |
|---|---|---|---|
| Habitat | - | - | - |
| Treatment group | - | - | - |
| Age | - | - | - |
| **Clutch size** | **32.572** | **1** | **<0.001** |
| Hatch date$^1$ | 0.677 | 1 | 0.411 |
| Hatch date$^2$ | 0.000 | 1 | 0.999 |
| **Habitat × treatment group × Age** | **19.404** | **3** | **<0.001** |

### b) Global model coefficients

| *Fixed Effects* | *Estimate* | *Standard Error* | *95% CI (Lower* | *Upper)* |
|---|---|---|---|---|
| **Intercept** (Control group in the urban habitat two days after hatching) | 6.439 | 0.493 | 5.437 | 7.237 |
| Treatment group: *Experimental* | -0.257 | 0.809 | -1.844 | 1.029 |
| Habitat: *Forest* | 0.858 | 0.690 | -0.539 | 2.086 |
| **Clutch Size** | **0.767** | **0.116** | **0.506** | **0.978** |
| Hatch date | -0.042 | 0.051 | -0.146 | 0.052 |
| Age: *day six* | -0.857 | 0.479 | -1.762 | 0.033 |
| **Age: *day 12*** | **-3.786** | **0.479** | **-4.791** | **-2.869** |
| **Age: *fledged*** | **-5.214** | **0.479** | **-6.036** | **-4.414** |
| Treatment group: *Experimental* × Habitat: *Forest* | -0.117 | 0.978 | -2.068 | 1.442 |
| Habitat: *Forest* × Age: *day six* | 0.607 | 0.655 | -0.662 | 1.698 |
| **Habitat: *Forest* × Age: *day 12*** | **3.411** | **0.655** | **2.13** | **4.449** |
| **Habitat: *Forest* × Age: *Fledged*** | **4.180** | **0.661** | **2.783** | **5.194** |
| Treatment group: *Experimental* × Age: *day six* | 0.011 | 0.689 | -1.311 | 1.217 |
| **Treatment group: *Experimental* × Age: *day 12*** | **2.709** | **0.689** | **1.442** | **3.761** |
| **Treatment group: *Experimental* × Age: *Fledged*** | **3.291** | **0.689** | **2.059** | **4.289** |
| Treatment group: *Experimental* × Habitat: *Forest* × Age: *day six* | -0.361 | 0.945 | -2.299 | 1.186 |
| **Treatment group: *Experimental* × Habitat: *Forest* × Age: *day 12*** | **-2.934** | **0.945** | **-4.869** | **-1.296** |
| **Treatment group: *Experimental* × Habitat: *Forest* × Age: *Fledged*** | **-3.452** | **0.954** | **-5.376** | **-1.521** |

***Random effects***

| *Groups* | *Variance* | *Standard deviation* |
|---|---|---|
| Clutch ID | 1.731 | 1.316 |
| Residual | 1.607 | 1.268 |

**Table S21. a)** Likelihood-ratio tests (LRT) for the following predictors explaining the proportion of nestlings alive: the treatment group × habitat × age interaction, clutch size, and hatch date. Clutch ID was included as a random effect. Significant P-values are highlighted in bold. "*df*" = degrees of freedom for the LRTs. Superscripts "1" and "2" refer to linear and quadratic terms, respectively. **b)** Model coefficients from the binomial generalised linear mixed model the effects of the following predictors on the proportion of nestlings alive: the treatment group × habitat × age interaction, number of hatched eggs, and hatch date. The standard errors and 95% confidence intervals are provided for each model coefficient. N=60 broods.

| Proportion of nestlings alive (Alive/Dead) | | | |
|---|---|---|---|
| **a) Likelihood-ratio test results** | | | |
| ***Predictors*** | $\chi^2$ | ***df*** | ***P-value*** |
| Habitat | - | - | - |
| Treatment group | - | - | - |
| Age | - | - | - |
| Hatch date[1] | 0.375 | 1 | 0.540 |
| Hatch date[2] | 0.061 | 1 | 0.805 |
| **Number of hatched eggs** | **8.154** | **2** | **0.017** |
| **Habitat × treatment group × Age** | **8.761** | **3** | **0.033** |

| **b) Model coefficients** | | | | |
|---|---|---|---|---|
| ***Fixed Effects*** | ***Estimate*** | ***Standard Error*** | ***95% CI (Lower \| Upper)*** | |
| **Intercept** (Control group in the urban habitat two days after hatching) | 2.462 | 0.526 | 1.413 | 3.520 |
| Treatment group: *Experimental* | 1.014 | 0.889 | -0.728 | 1.659 |
| Habitat: *Forest* | 0.081 | 0.794 | -1.528 | 2.840 |
| Hatch date[1] | -0.038 | 0.058 | 0.153 | 0.062 |
| **Number of hatched eggs** | **0.359** | **0.129** | **0.110** | **0.080** |
| Age: *day six* | -0.940 | 0.408 | -1.744 | -0.139 |
| **Age: *day 12*** | **-3.177** | **0.425** | **-4.025** | **-2.341** |
| **Age: *fledged*** | **-4.318** | **0.461** | **-5.240** | **-3.414** |
| Treatment group: *Experimental* × Habitat: *Forest* | 0.931 | 1.199 | -1.327 | 3.426 |
| Habitat: *Forest* × Age: *day six* | 0.542 | 0.607 | -0.646 | 1.736 |
| **Habitat: *Forest* × Age: *day 12*** | **2.613** | **0.612** | **1.423** | **3.826** |
| **Habitat: *Forest* × Age: *Fledged*** | **3.019** | **0.626** | **1.802** | **4.262** |
| Treatment group: *Experimental* × Age: *day six* | -0.567 | 0.688 | -1.327 | 0.762 |
| Treatment group: *Experimental* × Age: *day 12* | 1.349 | 0.695 | -0.017 | 2.696 |
| Treatment group: *Experimental* × Age: *Fledged* | 1.416 | 0.725 | -0.011 | 2.813 |
| Treatment group: *Experimental* × Habitat: *Forest* × Age: *day six* | -0.497 | 1.022 | -2.532 | 1.493 |
| **Treatment group: *Experimental* × Habitat: *Forest* × Age: *day 12*** | **-2.248** | **1.024** | **-4.295** | **-0.252** |
| **Treatment group: *Experimental* × Habitat: *Forest* × Age: *Fledged*** | **-2.499** | **1.035** | **-4.379** | **-0.294** |

| ***Random effects*** | | |
|---|---|---|
| ***Groups*** | ***Variance*** | ***Standard deviation*** |
| Clutch ID | 2.287 | 1.512 |
